# Interfacial residues in protein-protein complexes are in the eyes of the beholder

**DOI:** 10.1101/2023.04.24.538134

**Authors:** Jayadevan Parvathy, Arangasamy Yazhini, Narayanaswamy Srinivasan, Ramanathan Sowdhamini

**Affiliations:** Molecular Biophysics Unit (MBU), Indian Institute of Science, Bangalore; Interdisciplinary Mathematical Sciences Initiative (IMI), Indian Institute of Science Bangalore.; National Center for Biological Sciences (NCBS), Bangalore; Max Planck Institute for multidisciplinary sciences, Germany

**Author notes:** This article is dedicated to the memory of Prof. N. Srinivasan. **Funding Information** Parvathy’s PMRF grant and NS’s JC Bose fellowship.

**Keywords:** Interface residues, protein-protein interactions, Accessible Surface Area (ASA), interaction energy, hotspot residues, residue conservation.

## Abstract

Interactions between proteins are vital in almost all biological processes. The characterization of protein-protein interactions helps us understand the mechanistic basis of biological processes, thereby enabling the manipulation of proteins for biotechnological and clinical purposes. The interface residues of a protein-protein complex are assumed to have the following two properties: a) they always interact with a residue of a partner protein, which forms the basis for distance-based interface residue identification methods, and b) they are solvent-exposed in the isolated form of the protein and become buried in the complex form, which forms the basis for Accessible Surface Area (ASA)-based methods. The study interrogates this popular assumption by recognizing interface residues in protein-protein complexes through these two methods. The study shows that a few residues are identified uniquely by each method, and the extent of conservation, propensities and their contribution to the stability of protein-protein interaction varies substantially between these residues. The case study analyses showed that interface residues, unique to distance, participate in crucial interactions that hold the proteins together, whereas the interface residues unique to the ASA method have a potential role in the recognition, dynamics and specificity of the complex and can also be a hotspot. Overall, the study recommends applying both distance and ASA methods so that some interface residues missed by either method but crucial to the stability, recognition, dynamics and function of protein-protein complexes are identified in a complementary manner.

## 1. INTRODUCTION

Proteins play crucial roles in many biological processes like signalling, catalysis of metabolic processes, immune systems, and transporting molecules. To perform this wide range of functions, proteins interact with other biomolecules^1–3^. Therefore, the characterization of a protein-protein interface is vital in understanding their binding affinity, function etc., and is of utmost importance in their experimental mutagenesis studies, predicting protein-protein networks, designing drug targets, and engineering proteins. There are several experimental methods, like X-ray crystallography, Nuclear Magnetic Resonance (NMR) etc., for identifying the interface residues in a complex. However, these experimental methods are time-consuming, labour-intensive and have associated challenges like the protein not being amenable to experimental conditions for structure determination, difficulty in getting high-quality crystals and purification and expression of protein samples prone to aggregation. Hence various computational approaches for the same have become crucial. When the experimental structure of the complex is not available, homology-based methods or docking approaches are generally used to predict the structure of the complex and interface residues. Modeller is one such homology-based method that can also be employed to build protein-protein complex structures using known structures of homologous complexes^4^. HADDOCK and HDOCK, given the structure of the individual proteins, use an iterative algorithm that is energy and shape complementarity driven to predict the binding pose of two proteins in their complex form^5, 6^.

Recently developed Machine learning-based methods, namely AlphaFold2 and RoseTTaFold, have achieved unprecedented accuracy in protein structure prediction. AlphaFold2 is the first method to predict protein structures at atomic resolution, even without a homologous template structure. High accuracy is achieved by an advanced deep learning algorithm combined with various evolutionary, physical and geometrical constraints as features for training the model^7^. RoseTTaFold, which shows comparable performances with AlphaFold2, employs a 3-track network (instead of a 2-track network as in AlphaFold2) which integrates the sequence information, two-dimensional distance maps, and knowledge about the three-dimensional coordinates^8^. Both these methods have been extended to predict structures of protein-protein complexes in a way that bypasses the traditional approach of predicting the models of individual proteins and then docking them to get a complex structure. The network used in RoseTTaFold, enables rapid generation of accurate protein-protein complex models from sequence information alone, as the network can easily handle chain breaks in a single chain^8^. AlphaFold-Multimer is an algorithm that utilizes the protocol of AlphaFold2 by explicitly training it for multimeric inputs of known stoichiometry^9^. This helps the accurate prediction of protein complexes, especially for homomeric complexes. The most recent “FoldDock” is another extension of the AlphaFold2 algorithm incorporating optimized multiple sequence alignments, which folds and docks the heterodimeric protein-protein pairs simultaneously with an accuracy close to that of AlphaFold-Multimer^10^.

Several other machine learning-based methods classify a residue pair as interacting or not, provided some features associated with the residue pair are known. These algorithms are rapid and reported to have substantial accuracy in predicting interface residues^11^. There are sequence-based methods which use features like amino acid type and Position Specific Scoring Matrices (PSSM). At the same time, structure-based machine learning methods use features like Relative Accessibility Surface Area (RASA) and depth index to predict the interface residues. For example, PPiPP is a sequence-based method that uses a neural network for prediction^12^. In contrast, PAIRPRED uses both sequence and structure-based features and a Support Vector Machine (SVM) algorithm for interface prediction^13^. Many supervised machine learning-based tools use either distance-based or ASA-based criteria to label a residue pair as interacting. For example, in PAIRPRED, two residues belonging to two different proteins in a complex are considered to be interacting if the distance between any two heavy atoms of those residues in the bound conformations of their proteins is less than or equal to 6.0 Å^13^. Alternatively, in several other methods, a residue is labelled an interfacial residue if the change in its Accessible Surface Area (ASA) upon complexation is larger than 1 Å^2^ ^11, 14^.

The interface residues in a given protein-protein complex are identified using two widely used methods: a) distance-based and b) ASA–based^15–19^. In distance-based methods, if any of the atoms in that residue pair come closer than a distance cut-off chosen, the residue pair is considered interacting. This method generally captures the residues in the interface that contributes to the stability of the complex through direct interactions with residues in its partner protein. However, the protein-protein interface additionally comprises other residues that aid the stability of the complex by being buried from the solvent or through the intra-protein interactions they form. An ASA-based method picks residues that are exposed when the protein is in its isolated form and becomes buried upon binding with its partner protein as interfacial residues^20^. To the best of our knowledge, no study has focused on the interface residues unique to these methods.

In general, an interface residue in a protein-protein complex is assumed to always physically interact with a residue in its partner protein through stabilizing interactions and to be placed at the geometrical interface. According to the classical definition, the residues at the geometrical interface are considered to be solvent-exposed in the unbound form of the protein to which it belongs and become buried upon complex formation^19, 20^. However, it remains unclear whether interface residues possess both characteristics, i.e. making physical interactions and being placed at the geometrical interface. Generally, either the distance-based or ASA-based method is used to identify interface residues. Using only one of the methods might disregard some of the residues with crucial roles at the interface. In this study, we aimed to study interface residues with respect to the physical interactions and appropriate geometrical placement. From the analysis of 1153 crystal structures of protein-protein complexes, a considerable proportion of residues was found to be holding only one of the two characteristics. Detailed comparative studies on interface residues identified uniquely by distance-based or ASA-based methods were performed. The sequence conservation pattern and residue-wise energy analyses revealed that interface residues uniquely identified by distance methods have different attributes compared to interface residues uniquely identified by the ASA method. Further detailed case studies demonstrate different roles of uniquely identified interface residues in protein-protein interactions.

## 2. MATERIALS AND METHODS

### 2.1 Dataset preparation

The dataset was prepared using the crystal structures of protein-protein complexes retrieved from the Protein Data Bank (PDB)^21^. We screened the total crystal structures of 1,61,965 entries deposited in PDB (April 2022) with the following criteria: i) resolution better than 2Å, ii) only protein molecules in the structure, iii) only two chains in the structure, iv) no missing residues in the non-terminal regions, v) no missing backbone atoms, vi) protein length greater than or equal to 30 amino acids and vii) Rfree value less than or equal to 30%. The resultant 1195 crystal structures were repaired using the “RepairPDB” module in FoldX to remove energetically unfavourable interactions, incorrect torsion angles and Van der Waals clashes^22^. There were short contacts in structures even after the repair. Hence, Molprobity was used to recognize and remove such entries in the dataset with steric clashes^23^. In Molprobity, two atoms with an overlap of more than 0.4Å are considered clashing. The ‘clashscore’ reported in Molprobity is the number of severe clashes per 1000 atoms. Percentile scores are given for clash score relative to the group of PDB structures within 0.25 Å of the input structure’s resolution. A cut-off of 66 was recommended by Molprobity. Hence, 42 entries in the dataset with a clash score percentile < 66 were removed, culminating in a final dataset consisting of 1153 crystal structures (Figure 1A).

**FIGURE 1.**
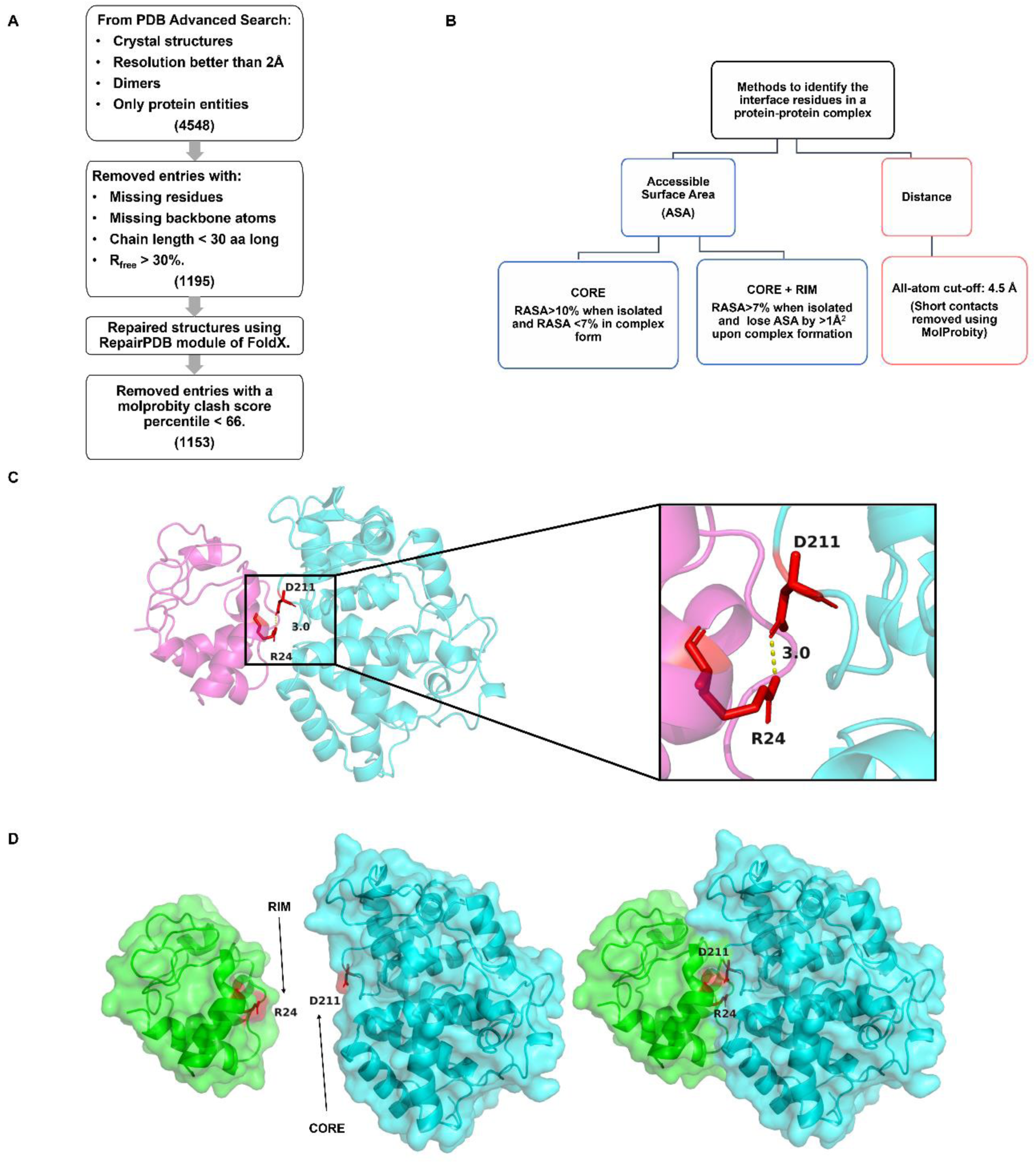
A) Flowchart of dataset generation. (B) Distance and ASA-based criteria that were used to identify interface residues in protein-protein complexes in the dataset. Cartoon visualization of interface residues identified by (C) distance method, (D) core and rim residues identified by ASA method, generated by Pymol (The PyMOL Molecular Graphics System, Version 2.5 Schrödinger, LLC).

### 2.2 Identification of interface residues by distance method

In distance-based methods, many criteria can be used to identify interacting residues in protein-protein complexes. For example, two residues can be accounted as interacting if the distance between their C^α^ atoms is less than or equal to 9 Å or if the distance between any two atoms is less than or equal to their corresponding van der Waals radii plus 0.5^15^. Similar criteria concerning C^β^ atoms or considering all atoms in the residue pair have also been used to identify interacting residues^16^. In this study, two residues are considered to interact with each other if the distance between any two atoms in the residue pair is less than or equal to 4.5 Å (Figure 1B)^24^. Residues in the interface that stabilize the complex by forming direct contact with the residues in its partner are hence captured. Using an in-house Python script and a Python package called Biopython, such interacting residue pairs in the whole dataset of 1153 complexes were identified^25^. Figure 1C shows an example of an interacting residue pair in the crystal structure of the Leishmania major peroxidase-cytochrome c complex (PDB code: 4GED), present in the dataset used in the study. One of the atoms of the Asp residue in chain A and Arg from chain B is placed at a distance of 3 Å, and hence these residues are considered to be interacting.

The intra-protein interactions in the complexes were identified using the same distance criteria. The interacting pair of residues (both inter-protein and intra-protein) are removed from the analysis if they have short contacts (excluding the short contacts involving Hydrogens) identified by Molprobity. The type of inter-protein and intra-protein interactions were identified using a web server Proteins Interactions Calculator (PIC)^26^. Some of the analyses in the following sections were performed using three other distance criteria to validate that the observations are independent of the distance criterion that is adopted in the study. Other distance criteria include C^α^ – based and C^β^ – based distance thresholds. In one of the criteria, two residues are said to interact if their C^α^ atoms fall within a distance of 6.5 Å^27, 28^. In the other two criteria, a residue pair is accounted to be interacting if the distance between their C^β^ atoms is within a distance of 7 Å^29^ and 4.5 Å^30, 31^, respectively.

### 2.3 Identification of interface residues by ASA method

In ASA-based methods, those residues in a protein-protein complex that are solvent-exposed in the unbound form of the protein to which it belongs and become buried upon binding with the partner protein are considered to be at the geometrical interface. This study identifies two categories of interface residues using the ASA-based method^17, 18^. A residue that is well solvent-exposed in the isolated form of the protein, as reflected by the Relative Accessibility Surface Area (RASA) value of greater than 10% and becomes buried in the complex form with a RASA value less than 7%, is located at the centre of the interface and categorized as ‘core’ residues. The residues partially exposed in the unbound form of the protein (RASA> 7%) and lose the ASA by more than 1 Å^2^ include residues at the centre as well as the periphery of the interface. Those residues that are placed at the periphery of the interface are named ‘rim’ residues. This method identified a more extensive set of residues that are part of the geometrical interface and lie at the centre as well as the periphery of the interface (Figure 1B). Naccess^32^ has been used to calculate residue-wise ASA and RASA values in the complex and isolated forms of the protein. It is a freely available software that enables the accessibility surface area calculations of proteins and nucleic acids. A Python script was used to identify the core and rim (Figure 1D) residues based on the values generated by naccess. In Figure 1D, the Asp residue has a RASA value of 62.6% in the unbound form and 5.8% in its bound form; hence it is a core residue. Whereas the Arg residue has a RASA of 44.2% in its unbound form and 9.9% in its bound form, and its ASA value reduces from 105.55 Å^2^ to 23.66 Å^2^ upon complex formation. Therefore, it is a rim residue and not a core residue.

The analysis focuses on the interface residues that are unique to each method which excludes the residues that are commonly identified by both methods. The interface residues identified solely by the distance-based method will be referred to as ‘unique to distance’ and solely by the ASA-based method as ‘unique to ASA’ interface residues in the following sections.

### 2.4 Percentage of interface residues uniquely identified by distance and ASA methods

The percentage of the total interface residues identified uniquely by each method was calculated. The total number of interface residues unique to a method for each complex in the dataset was calculated using a Python script. The count was normalized by the total number of interface residues identified in the protein-protein complex to account for the variation in the number of residues identified by each method. The average values of these percentages for the whole dataset were then compared between distance and ASA methods. The formulae used are as follows:

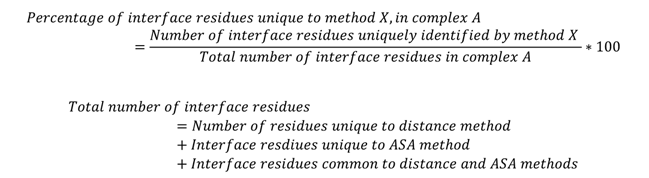

### 2.5 Propensity calculations

We probed to identify the nature of residues with a higher tendency to be interface residues unique to each method. The Chou-Fasman propensity^33^ calculations were employed to measure them as follows:

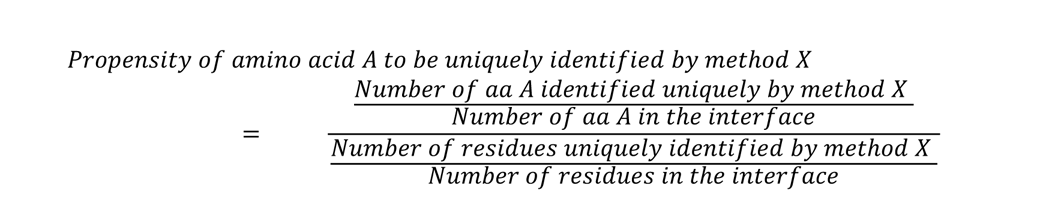

### 2.6 Calculation of residue conservation scores

ConSurf was used to calculate conservation scores for each interface residue unique to distance and ASA methods^34^. Given an input structure, the ConSurf algorithm extracts the corresponding sequence, identifies the homologs and finds the representatives using the clustering by CD-HIT^35^. From the multiple sequence alignments and the phylogenetic tree constructed for the identified representative homologs, the evolutionary rates are calculated using a Bayesian statistics-based algorithm called Rate4Site^36^. The HMMER method was used to search against the UNIREF90 database for one iteration and an E-value cut-off of 0.0001 to search for homologues in the context of ConSurf. MAFFT-LINSI was employed for deriving multiple sequence alignments for the identified representative homologs. The ConSurf grades obtained in the final output are the evolutionary rates of each residue position normalized to values ranging from 1 to 9. A grade of 9 indicates high conservation, a slow evolution at the given residue position and hence its importance in the structure or function of the protein. Generally, a residue with a ConSurf grade greater than six is considered partially conserved or conservatively substituted residue, and a grade less than or equal to six is considered to be poorly conserved. Homologs could not be detected for nine complexes in the dataset; hence, the conservation analyses were conducted for the dataset, excluding these nine entries. ConSurf gives another parameter called “amino acid variety”, along with the evolutionary scores. For each residue, this indicates the different amino acid residues that occur at that position in the multiple sequence alignment generated by ConSurf, along with the percentage of occurance. Figure S1 shows the representation of this parameter by sequence logos generated using WebLogo^37^. For each residue position the sequence logo shows the different amino acid residue type that occurs at that position in the input multiple sequence alignment. The size of the logo represents the frequency of the residue type in the MSA.

### 2.7 Calculation of inter-residue interaction energies

A software called PPCheck^38^ was used to measure the contribution of each interface residue (unique to distance and the ASA method) to the overall interaction energy of the complex. Given a structure with two interacting partners, PPCheck calculates the interchain energies: namely, hydrogen bonding, interchain Van der Waals interaction (hydrophobic interactions) and interchain electrostatics (Figure S2). The input to PPCheck in this study is the interface residue unique to each method and its corresponding intra-protein partner in the case of the ASA method and both inter-protein and intra-protein partners in the case of the distance method. This is performed to calculate the residue-wise energetics in such a way that the local environment of the input residue is preserved^39^.

### 2.8 Identification of hotspot residues

Hotspot residues are specific residues at the interface of the protein-protein complex which contribute substantially to the stability of the complex. The mutation of hotspot residues will destabilize the complex. In computation-based methods, hotspot residues are defined as those residues in a complex that, when mutated to Alanine, lead to a ΔΔG > 2 kcal/mol^40^. The AlaScan module of FoldX has been used to determine such residues in each complex in the dataset^22^. The percentage of hotspots among the interface residues unique to distance and ASA methods were calculated as follows:

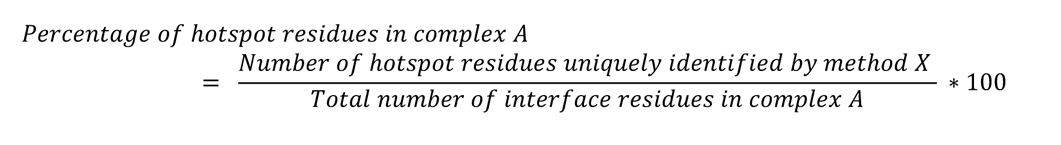

The percentage of hotspot residues among the residues unique to each method was calculated for each of the complexes, and the average values of each method for the whole dataset were compared.

### 2.9 Dataset imbalance

In the final dataset, the interface residues unique to ASA were more than double the interface residues unique to the distance method when the chosen distance and ASA-based criteria were implemented. This leads to dataset imbalance in certain comparative analyses in the case of conservation scores and residue-wise energies. A random selection was implemented using the ‘random’ module in Python to overcome the dataset imbalance. This module aids in sampling residues randomly from the residues unique to ASA and divides them into two equal halves, which will be referred to as “ASA_SET1” and “ASA_SET2” in the coming sections. In the conservation and interaction energy analyses, apart from the comparison between ASA and distance methods, the corresponding values for ASA_SET1 and ASA_SET2 are also compared with the values corresponding to the distance method.

### 2.10 Statistical significance test

A non-parametric test called “Mann-Whitney” at a 0.05 significance level was used to test the statistical significance of the difference in the distribution of conservation scores and residue-wise energy values corresponding to each residue type.

### 2.11 In silico mutagenesis

The position-specific *in silico* mutation for the case studies was performed using the PositionScan module of FoldX^22^. PositionScan mutates one amino acid to the remaining 19 amino acids and repairs the neighbouring residues. The mutated complexes from PositionScan were side-chain optimized using SCWRL4, a highly accurate algorithm for predicting the side-chain conformations^41^.

## 3. RESULTS AND DISCUSSION

### 3.1 ASA method tends to identify more unique interface residues

In the final dataset, 5832 interface residues were found to be unique to the distance method (6.02% of total interface residues in the dataset). In comparison, the interface residues unique to ASA are 12540 in number (12.95% of total interface residues in the dataset), which is more than double the interface residues unique to the distance method. This dataset imbalance is overcome using random selection. It was also observed that more than 75% of the interface residues unique to distance have a RASA value of less than or equal to 10%, in both the bound and isolated forms of the protein (Figure S3). This indicates that an interface residue happens to be unique to distance majorly because they tend to stay buried from solvent both in the bound and isolated forms of the protein.

From the examination of 1153 crystal structures in the dataset, the average values of the percentage of interface residues unique to distance and ASA methods were found to be 5.9% and 14.5%, respectively (Figure 2A). This is anticipated because the ASA method includes both core and rim residues corresponding to the centre and periphery of the interface. Interestingly, the same trend was observed upon using three other distance criteria, which are C^α^ and C^β^ -based. In the case of the ASA method, average values of 60.6%, 28.9%, and 83.1%, and in the case of the distance method, average values of 1.7%, 4.5%, and 0.2% were obtained while using a C^α^ -based cut-off of 6.5 Å (Figure S4A), C^β^-based distance cut-off of 7 Å (Figure S4C), and 4.5 Å (Figure S4E), respectively. This added confidence to the observation that the ASA method identifies more unique interface residues.

**FIGURE 2.**
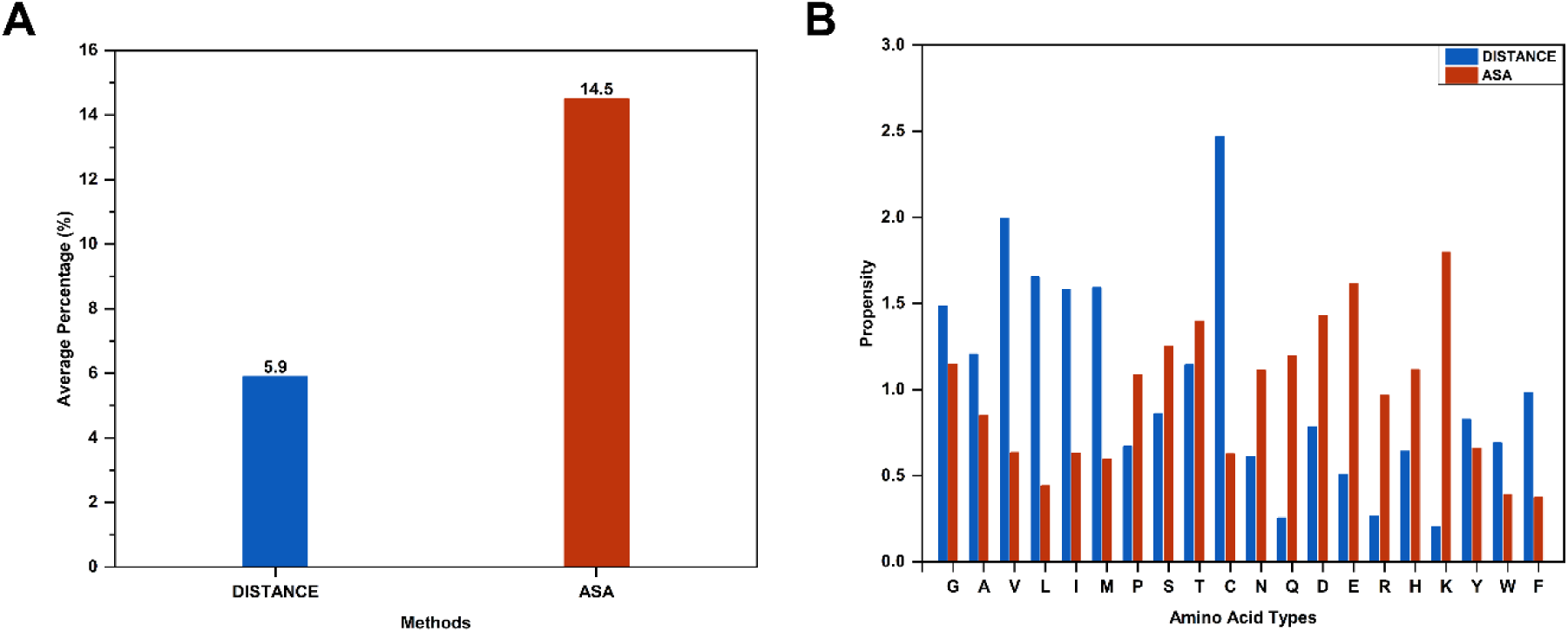
(A) Average percentages of interface residues uniquely identified by distance and ASA method. (B) Chou-Fasman propensity calculations to find the propensity of each amino acid type to be captured as unique to distance and ASA methods.

### 3.2 The propensity of interface residues unique to distance shows a contrasting trend with the residues unique to the ASA method

The propensity values of various amino acid residues to be unique to each method (Figure 2B) were examined. Apart from Gly, the propensity values to be unique to the distance method for hydrophobic residues like Ala, Val, Leu, Ile, and Met are greater than one and higher compared to the corresponding values for the ASA method. This trend is anticipated as hydrophobic patches are generally found in the protein-protein interaction interface, and they form an extensive network of inter-protein interactions. Also, from the detailed case studies and the dataset analyses, it was observed that the residues unique to the distance method failed to be captured by the ASA method because they remained buried in both the bound and isolated forms of the protein (Table S1 and Figure S3). Such residues are expected to be non-polar in nature. The same trend is seen in the case of Cys residues, and this is possibly due to the disulphides formed at the interface. The propensity for the ASA method shows an opposite trend compared to the distance method. The propensity values to be unique to the ASA method are greater than one and higher than the distance method for polar but uncharged residues, namely Thr, Ser, Asn, Gln and charged residues, namely Glu, Asp, Lys, and His.

Further, it was verified if these observations hold valid upon using other distance criteria. Upon using the C^α^ -C^α^ distance cut-off of 6.5 Å (Figure S4B), the propensity values to be unique to the distance method for Gly, Ala, Val, Ile, and Cys are still greater than one and higher compared to ASA values. The results are consistent in the case of ASA values for Asn, Gln, Glu, His and Lys residues. While considering the C^β^ - C^β^ distance cut-off of 7 Å (Figure S4D), the propensity values to be unique to the distance method for Gly, Ala, Val, Ile, and Cys are consistently greater than one and higher compared to ASA values. For Asn, Gln, Asp, Glu, His and Lys residues, the results remain consistent in the case of the ASA method. Similarly, while employing a C^β^ - C^β^ distance cut-off of 4.5 Å (Figure S4F), the propensity values to be unique to the distance method for Gly, Ala, and Leu are still greater than one and higher compared to ASA values. These results are consistent in the case of ASA values for Asn, Gln, Asp, Glu, His and Lys residues. The consolidated results from all four distance cut-offs showed that Gly and Ala residues consistently have propensity values greater than one and values greater than those corresponding to the ASA method. Similarly, in the case of the ASA method, Asn, Gln, Glu, His and Lys consistently show propensities greater than one and greater than that of the distance method consistently.

### 3.3 Interface residues unique to distance tend to be highly conserved

Generally, the interface residues crucial for the complex formation are highly evolutionarily conserved. Hence, we set out to look for if there is some conservation pattern specifically for the interface residues unique to distance and ASA methods. The percentages of interface residues unique to distance and ASA methods belonging to each ConSurf grade ranging from 1 to 9 are shown in Figure 3A. Interestingly, the distance method has higher percentages for ConSurf grades greater than six (16.4%, 18%, and 29.2% for ConSurf grades 7, 8, and 9, respectively) compared to residues unique to the ASA method (11.8%, 10.1%, and 11.2% for grades 7, 8, and 9 respectively). This suggests that the residues unique to the distance method tend to be highly conserved compared to the ones unique to the ASA method. Also, the trends for residues unique to the core and to the distance method are similar for almost all classes (Figure 3A). This is expected since the residues unique to distance form crucial hydrophobic or ionic interactions with the partner protein, and the residues unique to the core are located at the centre of the interface and are expected to be evolutionarily highly conserved. Similarly, the interface residues unique to the ASA method and those unique to specifically rim have similar percentages in all classes (Figure 3A). This shows that the residues unique to ASA tend to be less conserved compared to the distance method majorly due to the contribution from rim residues rather than the core residues.

**FIGURE 3.**
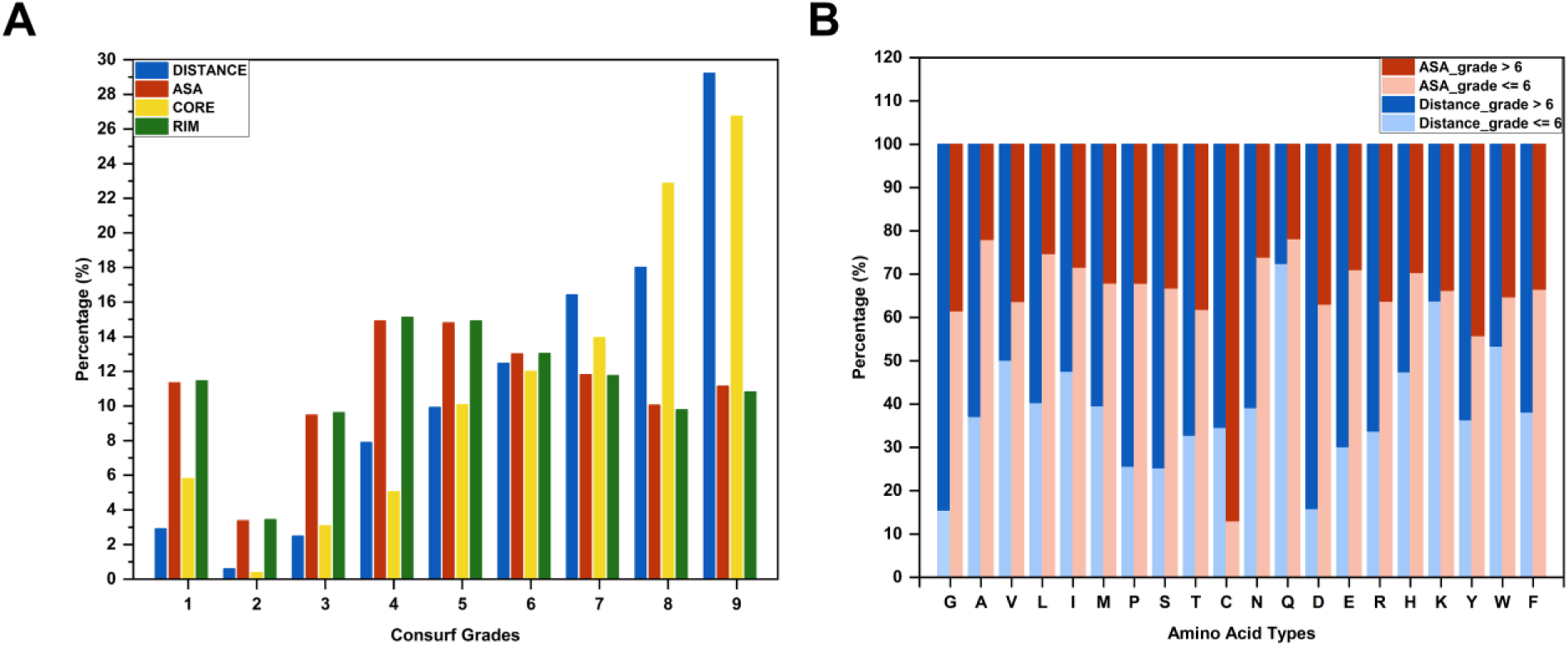
(A) Percentage corresponding to conservation score of residues unique to distance and ASA methods as well as core and rim residues identified from the ASA method. (B) Percentage of interface residues unique to distance method and ASA method with ConSurf grade <=6 and > 6, corresponding to each residue type.

The observation that the distance method predominantly identifies highly conserved residues as unique could be biased due to the imbalance in the dataset. Hence, to validate the observation, the percentages for residues unique to the distance method were compared with the values in ASA_SET1 and ASA_SET2 generated using the random selection (Figure S5A). The values for ASA, ASA_SET1 and ASA_SET2 were found to be similar for each ConSurf grade. This clearly validates the observation that interface residues unique to the distance method are highly conserved compared to that of the ASA method.

Next, the distribution of ConSurf grades for each of the 20 residue types was analyzed. Figure 3B shows the percentage of each amino acid with a ConSurf grade belonging to the two classes; (a) ConSurf grade greater than six, and (b) ConSurf grade less than or equal to 6. In the case of the distance method, most residues have a higher chance of belonging to the class with a ConSurf grade greater than six compared to less than or equal to 6, and corresponding percentage values are greater than 50. The exceptions are Val, Gln, Lys and Trp (49.9%, 27.5%, 36.2%, and 46.6%, respectively, for grades greater than six, and 50.1%, 72.5%, 63.8%, and 53.4%, respectively for grades less than or equal to 6). In the case of the ASA method, all residues except Cys have a higher tendency to be in the class of ConSurf grade less than or equal to 6 (13.1%) compared to the class of grade greater than 6 (86.9%). All the corresponding percentages are above 50. Hence, the results are in agreement with the observation that the interface residues unique to the distance method are predominantly conserved in nature, except for few amino acid residue types, namely Val, Gln, Lys, Trp, and Cys. While comparing the overall distribution of ConSurf grades for each residue type in the interface residue unique to distance versus ASA methods (Figure S5B) and distance versus rim (Figure S5D), the distributions are significantly different except for Lys residue (p-values 0.89 and 0.88, respectively). All the distributions cover the ConSurf grade range of 1 to 9. The mean and median of the distribution corresponding to the distance method lie at a range greater than six except for Val, Gln and Lys, whereas in the case of ASA and RIM, they lie below a ConSurf grade of 7. When ConSurf grades for core residues are compared with that of the distance method (Figure S5C), the distributions are significantly different only for Ala, Val, Leu, Met and Tyr residues (p-values 1.62 x 10^-4^, 0.03, 0.02, 0.01, and 0.04, respectively). Among these, except for Lys and Tyr, distributions corresponding to other residues have higher mean and median than the core and lie above 6. Overall, the observations from various analyses show that interface residues unique to distance are more conserved compared to the residues unique to ASA. The residues unique to ASA are observed to be conserved, provided they are at the core region of the interface.

### 3.4 Interface residues unique to the distance method contribute more favourably to the overall interaction energy of the complex

To examine the difference in the roles of interface residues unique to ASA and distance methods towards the overall stability of the complex, the residue-wise energy values from PPCheck were analyzed. The percentage of interface residues unique to each method belonging to various energy bins of length 5kJ/mol ranging from -20 kJ/mol to 30 kJ/mol was analyzed (Figure 4A). 95.6% of the interface residues unique to the distance method belong to the bin with an energy value of 0 to -5 kJ/mol, indicating favourable interaction. In contrast, only less than 8.2% of the interface residues unique to ASA have favourable interaction energy values. More than 90% of the interface residues unique to ASA have energy values in the positive range. The energy values for distance with ASA_SET1 and ASA_SET2 (Figures S6A and S6B) and energy values of distance with core (Figure 4B) and rim (Figure 4C) energy values were further compared. In all the comparisons, a higher percentage of interface residues unique to the distance method possess highly favourable energy values compared to ASA residues.

**FIGURE 4.**
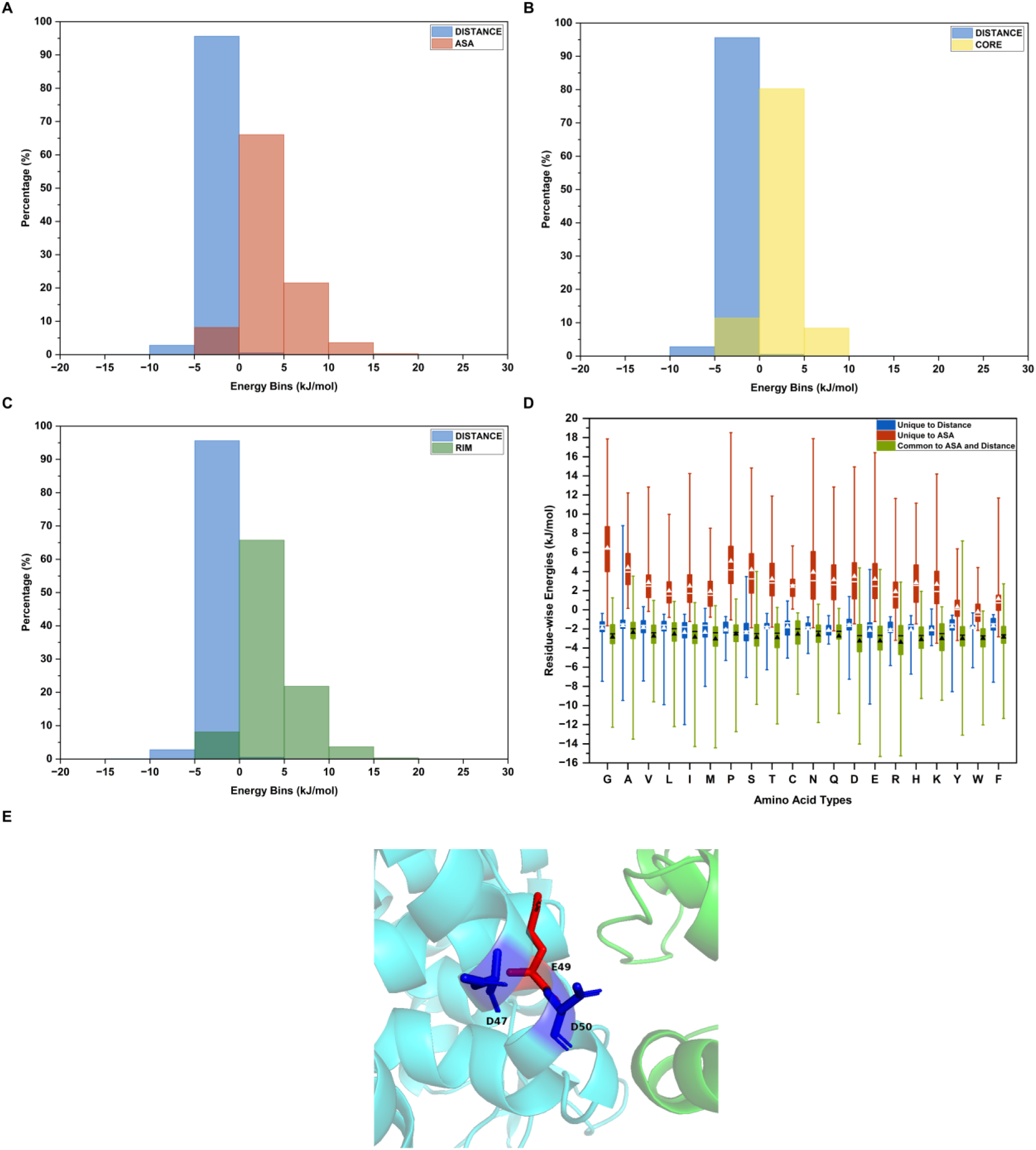
Histogram of residue-wise energies (kJ/mol) of interface residues compared between interface residues uniquely identified by (A) distance and ASA method, (B) distance and core residues in ASA method and (C) distance and rim residues in ASA method. X-axis has the residue-wise energies calculated using PPCheck^38^, binned with each bin having a 5 kJ/mol width. (D) Box plot showing the distribution of residue-wise energies of interface residues unique to distance, ASA and interface residues common to both methods for each amino acid residue type. The white star and line on the plot indicate the mean and median of the distributions, respectively. (E) Cartoon visualization showing interface residue unique to the rim (49 Glutamate, shown in red sticks) having a positive residue-wise energy due to repulsion from a similar charge in its surrounding (47 Aspartate and 50 Aspartate, shown in blue sticks) (PDB code: 4GED).

While examining the distribution of energy values for each residue type, the values corresponding to Cys residue were extreme compared to that of other residues. Due to the potential role of disulphide bridges in this, the energy values for each residue type for 686 complexes in the dataset without inter-protein and intra-protein disulphide bridges were analyzed. This analysis also holds that the energy values for most of the interface residues unique to ASA lie in the unfavourable region compared to distance. In the case of the distance method, most of the residues have energy values in the negative range except for a few outliers, which are positive (Figure 4D). Still, it was intriguing to see that most of the interface residues unique to the ASA method always have unfavourable residue-wise energy values. Hence, to confirm this observation, roughly 20% of the interface residues common to ASA and distance methods which is 9921 in number, were selected randomly, and the corresponding energy values were analyzed. Figure 4D shows the distribution of energy values for interface residues common to both methods apart from the unique ones. The distribution of energy values is mostly in the favourable energy range, apart from a few outliers. While comparing the energy values of each residue type that are unique to distance with the values corresponding to residues unique to the core and rim (Figure S6C and S6D), the same observation holds. Hence, it was concluded that the interface residues unique to the distance method contribute favourably to the overall interaction energy of the complex compared to those unique to the ASA method. The strong favourable energy values of residue unique to the distance method are possibly due to the formation of crucial stabilizing and evolutionarily conserved interacting ion pairs or hydrophobic patches at the centre of the interface. At the same time, the residues unique to ASA are highly polar or mostly charged in nature, as indicated by propensity calculations (Figure 2B). Thus, the highly unfavourable energies could be due to repulsion among these residues with the similarly charged proximal residues within the protein in the absence of the shielding effect by the solvent. For example, Figure 4E shows the Glu residue in Leishmania Major Peroxidase, an interface residue unique to the rim. This residue makes repulsive intra-protein interactions with two neighbouring similarly charged Asp residues in Cytochrome c in the complex (PDB code: 4GED), which contribute to the unfavourable energy value. Such rim residues provide specificity to the protein to recognise its right partner protein.

### 3.5 Distance method predominantly captures more hotspot residues as unique to it

It is essential to understand which method predominantly captures the hotspot residues as unique to it since these residues play a dominant role in the stability of the complex. The average values of the percentage of interface residues that are hotspots and unique to each method were compared. The distance method had a slightly higher value (3.8%) compared to the ASA method (2.91%) (Figure 5). The same observation holds true when the average value for the distance method is compared with both core (0.11%) and rim (2.8%) residues (Figure 5). These results may vary with the other distance cut-offs (Results are not reported here). We also observed that more than 95% of the hotspot residues unique to distance have RASA values of less than or equal to 10% in both the bound and isolated forms of the protein. This indicates that these hotspot residues tend to stay buried from the solvent in both the bound and isolated forms of the protein (Figure S7). Also, from the propensity calculations, we observed that the residues unique to distance are non-polar in nature (Figure 2B, S4B, S4D, and S4F). This suggests the existence of the slightly buried and non-polar layer of hotspot residues beneath the primary layer of crucial hotspot residues. We term such hotspot residues unique to distance, which are also observed to be non-polar and less accessible to the solvent as “secondary shell hotspots”. These will be further studied in detail in the future.

**FIGURE 5.**
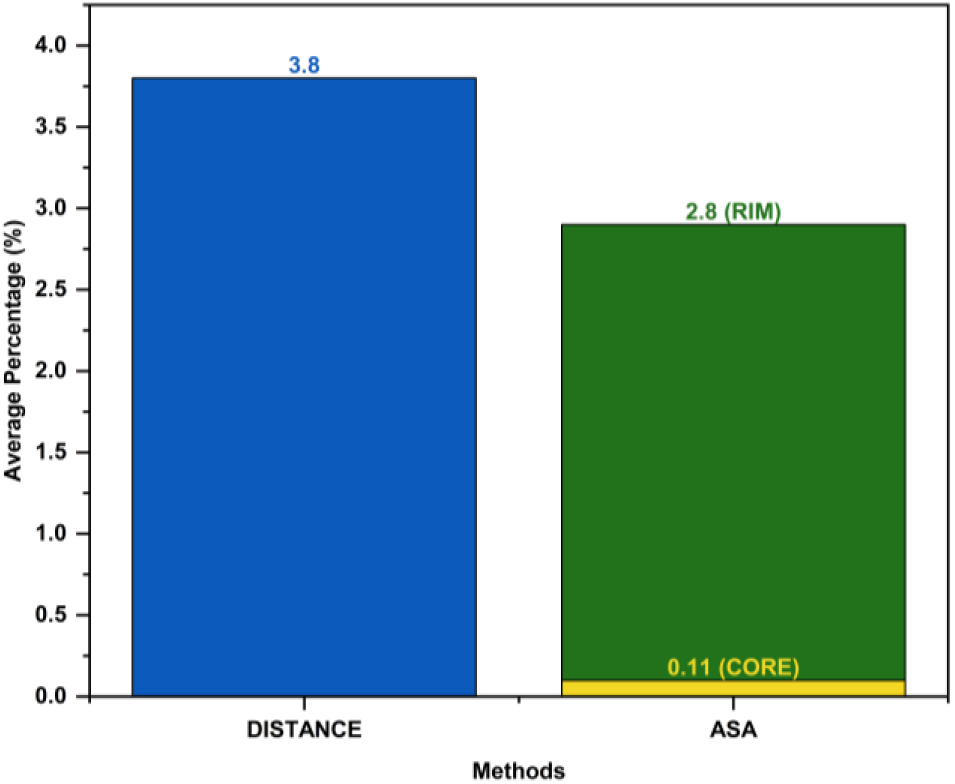
Average percentage of hotspot interface residues unique to distance, ASA methods, core and rim residues.

### 3.6 Case studies

To understand the biological relevance of interface residues unique to each method in a complex, crystal structures of four protein-protein complexes from the dataset were analyzed in detail, as discussed in the following sections.

#### 3.6.1 Case study-1

Vasohibins are tubulin tyrosine carboxypeptidases, the class of enzymes that releases the C-terminal Tyr from alpha-tubulin, converting tyrosine-terminated (Tyr) to detyrosinated (Glu) tubulin. The Small Vasohibin-Binding Protein (SVBP) has an allosteric role in the positive regulation of the detyrosination activity of Vasohibins^42, 43^. Crystal structures of the SVBP-bound form of the complex are available for two Vasohibins, VASH1 and VASH2. The sequence identity between VASH1 and VASH2 is roughly 52%. In VASH1-SVBP (PDB code: 6J7B), SVBP adopts an alpha-helical conformation and passes through a cavity formed by α1-α5 of VASH1, with its helix axis roughly parallel to that of α1. Specifically, residues 32–49 of SVBP make intermolecular contacts along the helix axis with VASH1 via two faces: 1) Interface 1, primarily involving hydrophobic interactions, and 2) Interface 2, consisting of electrostatic interactions (Figure 6A)^42^. In VASH2-SVBP (PDB code: 6J4P), the V2c (Predicted structured core domains of V2) structure can be subdivided into an N-and a C-terminal domain. The helical N-terminal domain (ND) wraps around a long helix formed by the C-terminal segment of SVBP to form a three-helix bundle. The C-terminal domain of V2c (CD) comprises the predicted catalytic residues of vasohibins. The ND and CD are packed together via SVBP, which is sandwiched between both domains. It has also been reported that the deletion of the ND disrupted the complex formation (Figure 7A)^43^. The crystal structures of the structured core domains of VASH1-SVBP heterodimer (PDB code: 6J7B)^42^ and that of VASH2 (V2c)-SVBP (PDB code: 6J4P)^43^, both in the presence of a Tyr residue-based inhibitor name epoY (A mimic of the enzyme-substrate α-tubulin), were analyzed in detail. They share a conserved catalytic triad (Cys169, His204 and Leu226 in VASH1 and Cys158, His193 and Leu215 in VASH2). Apart from the catalytic triad, loop 133QYNH136 is also conserved between VASH1 and VASH2. The loop 133QYNH136 aids in the allosteric role of SVBP by holding Tyr134, a residue important for peptide detyrosination, in a favourable position. Figure S8 shows the structure-based alignment of VASH1 and VASH2 with these conserved regions highlighted. It has been reported that SVBP binds to VASH1 and VASH2 with Kds of roughly 42 nM and 18 nM, respectively^42, 43^. Even though it is reported that these two complexes share all known motifs and display similar functions, they differ substantially in the strength of binding^44^. This motivated us to analyze the differences in the interface and, specifically, the role of residues unique to distance and ASA methods in the overall binding and specificity of these complexes.

**FIGURE 6.**
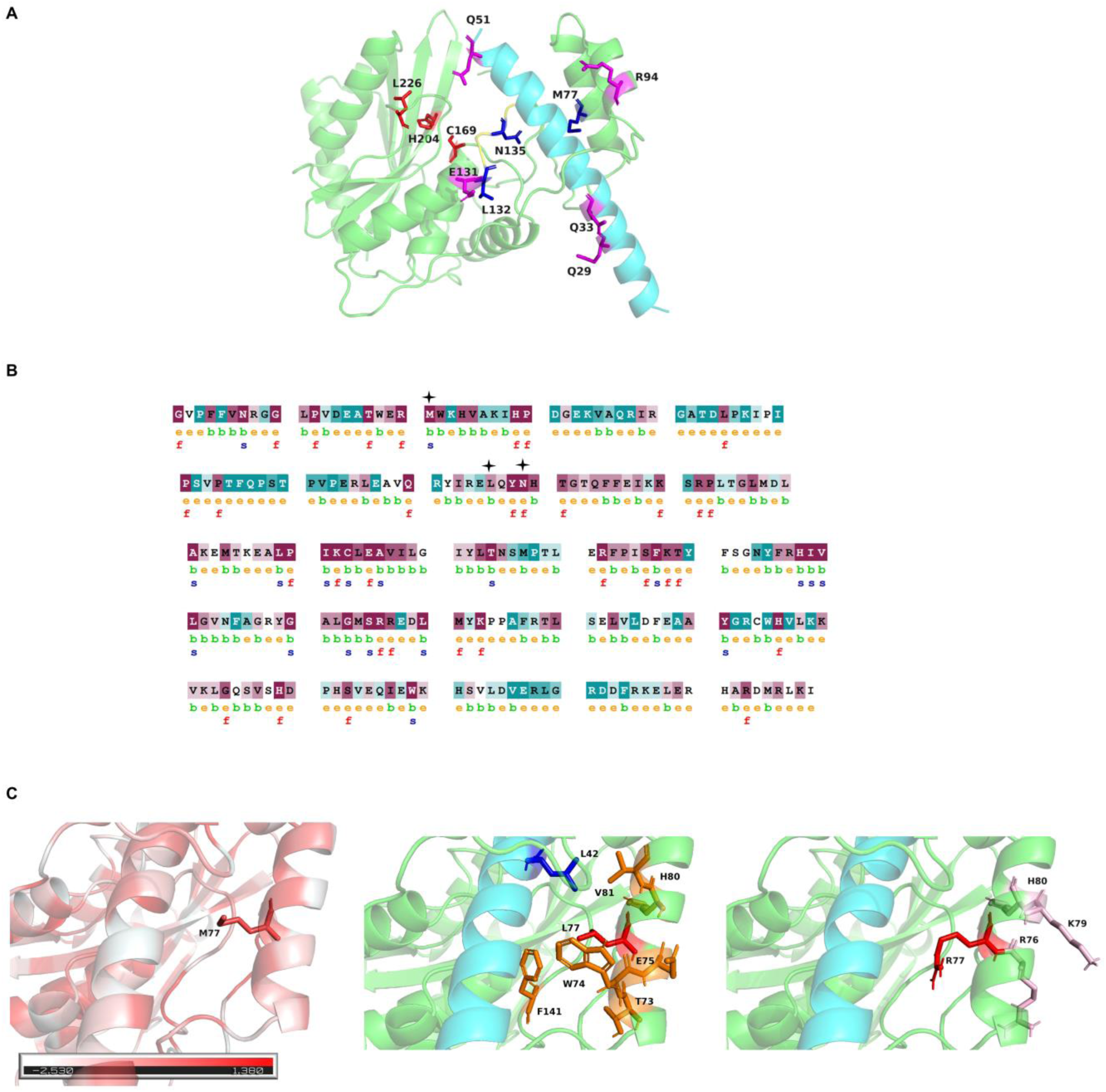
(A) Cartoon representation of VASH1 (green) and SVBP (cyan) complex structure (PDB code 6J7B). The catalytic triad (Cys169, His204 and Leu226 in VASH1) and the 133QYNH136 loop conserved between VASH1 and VASH2 are shown in red sticks and yellow cartoons, respectively. Interface residues unique to the core, rim and distance are shown in orange, magenta and blue sticks, respectively. (B) Amino acid sequences colour coded by conservation score for protein VASH1. The figures were generated using ConSurf^34^. The residues discussed in the case study are marked in black stars. (C) Cartoon representation of the wild-type VASH1 (green) -SVBP (cyan) complex structure (PDB code: 6J7B) (Left) coloured according to hydrophobicity values. Met77 of VASH1, a residue uniquely identified by the distance method in the VASH1-SVBP complex that is subjected to *in-silico* mutation, is shown in stick representation. The interaction network in the *in silico* mutant M77L and M77R are shown in the middle and the right panels. The inter-protein and intra-protein interaction partners and the spatially proximal residues (within a distance of 4.5 Å) that could cause charge-charge repulsions are shown in blue, orange, and pink sticks, respectively.

**FIGURE 7.**
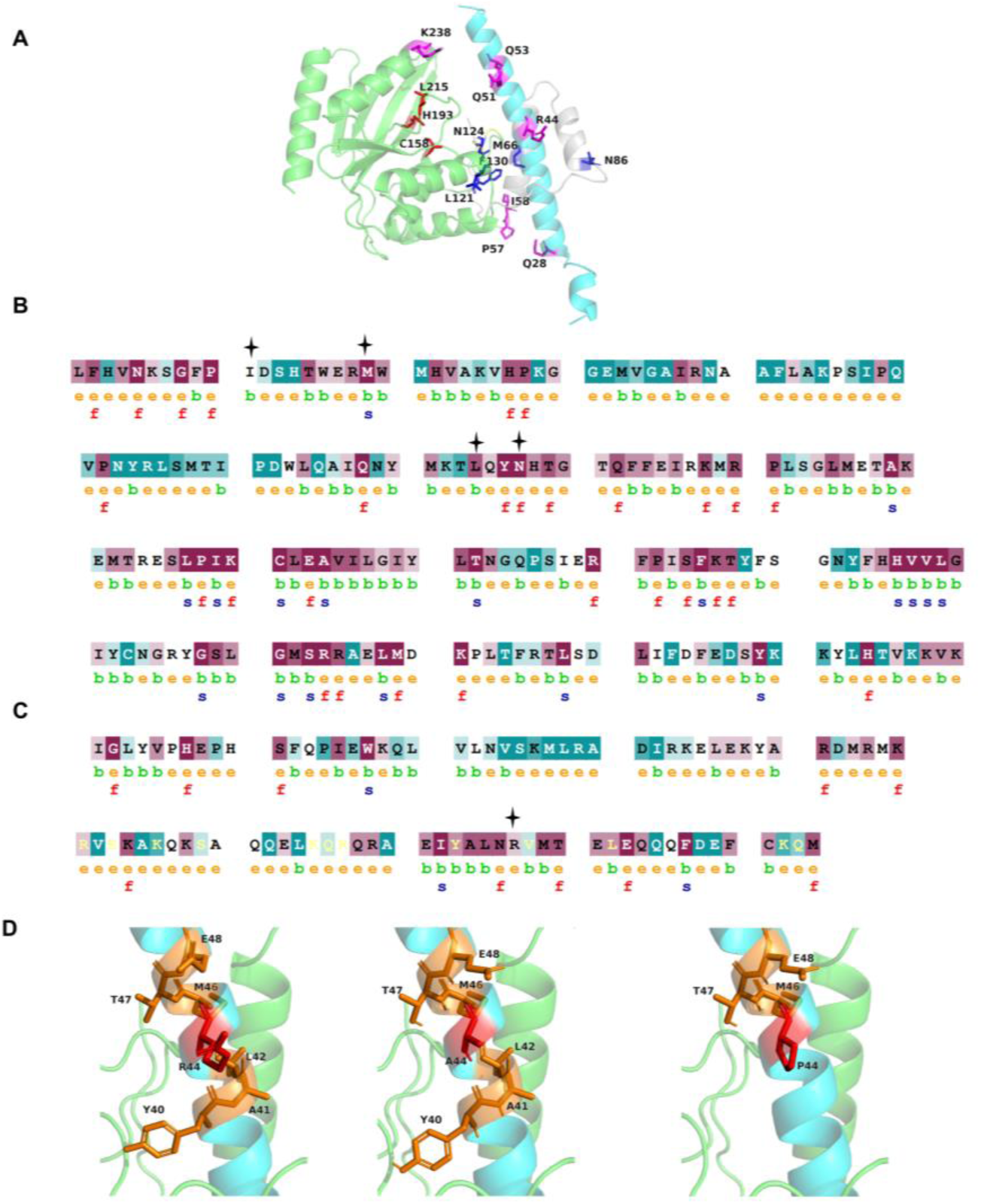
(A) Cartoon representation of VASH2 and SVBP (cyan) complex structure (PDB code 6J4P). The N-terminal domain (ND) and C-terminal domain of VASH2 are in grey, and green cartoons, respectively. The catalytic triad (Cys158, His193 and Leu215 in VASH2) and the 133QYNH136 loop conserved between VASH1 and VASH2 are shown in red sticks and yellow cartoons, respectively. Interface residues unique to the core, rim and distance are shown in orange, magenta and blue sticks, respectively. Amino acid sequences colour coded by conservation score for protein (B) VASH2 and (C) SVBP. The figures were generated using ConSurf^34^. The residues discussed in the case study are marked in black star. (D) The interaction network in the wild-type VASH2 (green) -SVBP (cyan) complex structure (PDB code: 6J4P), *in silico* mutant R44A and R44P, are shown in the left, middle and right panels, respectively. Arg44 of VASH1 that is uniquely identified by the ASA method in the VASH2-SVBP complex that is subjected to *in-silico* mutation, is shown in stick representation. The intra-protein interaction partners are shown in orange sticks.

#### 3.6.2 Case study-2

*Leishmania major* is the causative agent of leishmaniasis. The host produces reactive oxygen species like hydrogen peroxide as a protective mechanism. In response, *L. major* produces a heme peroxidase, L. Major Peroxidase (LmP), which helps the Lm from oxidative stress by catalyzing the peroxidation of its mitochondrial cytochrome c, LmCytC. The crystal structure (PDB code: 4GED) of LmP in complex with its substrate LmCytc at a resolution of 1.84 Å was analyzed in detail in this study (Figure 8A)^45^. The unbound crystal structures of both LmP (PDB code: 3RIV)^46^ and LmCytC (PDB code: 4DY9)^47^ were available while the study was conducted. It has been reported that the individual and complex structures do not have any major conformational changes apart from the minimal side chain reorientations at the interface upon complex formation^48^. RMSD from structural superposition by TM-Align^49^ for LmP in bound and unbound forms is 0.32Å, and that for LmCytC is 0.48Å. This enables comparing various interactions formed by these proteins in their bound and unbound forms.

**FIGURE 8.**
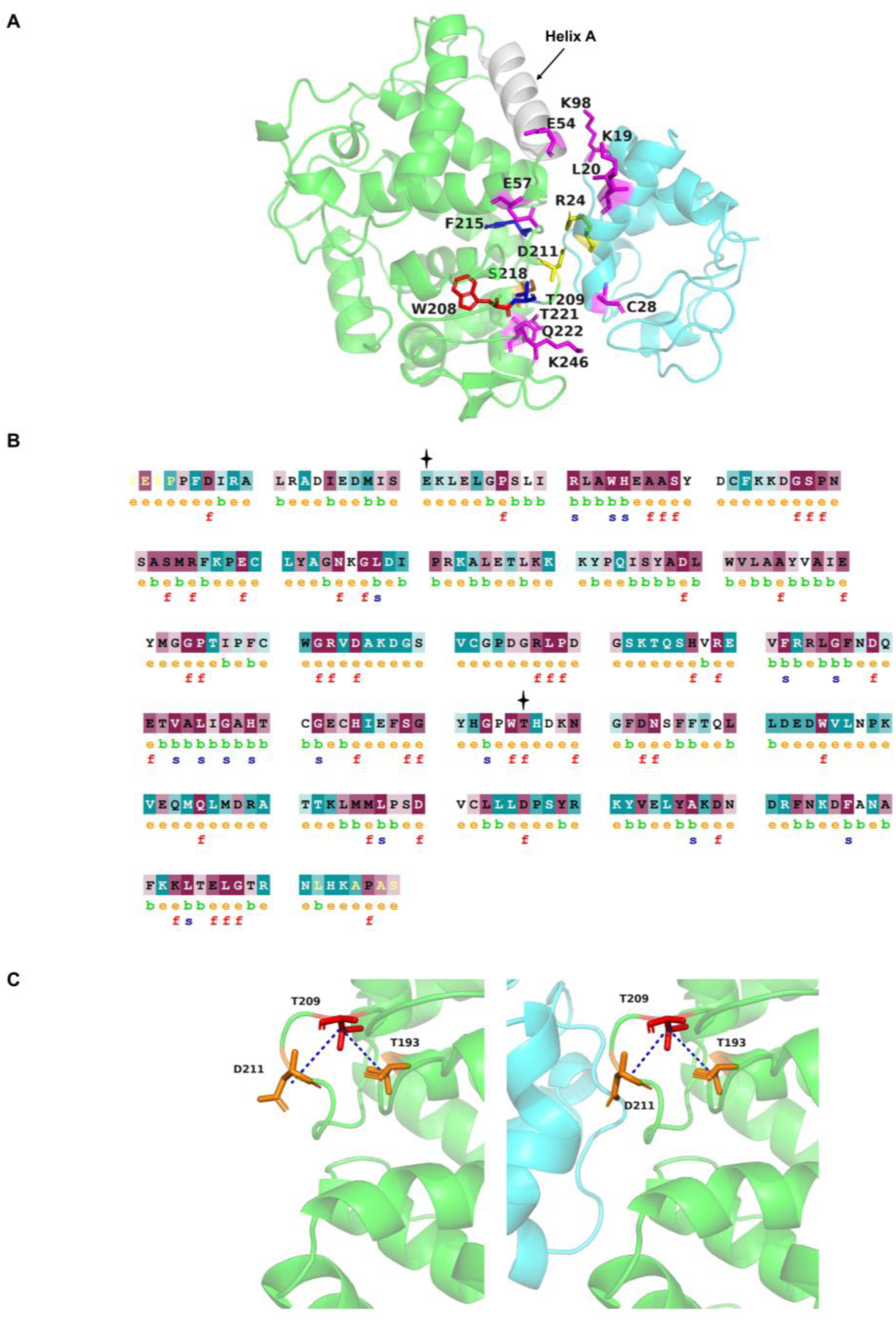
(A) Cartoon representation of LmP (green) and LmCytC (cyan) complex structure (PDB code: 4GED). The non-catalytic secondary binding site, helix A in the grey cartoon, the tryptophan radical Trp208 in red sticks, and the ion pair LmP Asp211 – LmCytC Arg24 crucial for complex formation and activity in yellow sticks. Interface residues unique to the core, rim and distance are shown in orange, magenta and blue sticks, respectively. (B) Amino acid sequences colour coded by conservation score for protein LmP. The figures were generated using ConSurf^34^. The residues discussed in the case study are marked in black star. (C) The interaction network of Thr209 (red sticks) of LmP in its unbound form (PDB code: 3RIV) (left panel) and bound form (Right panel) (PDB code: 4GED). The intra-protein interaction partners are shown in orange sticks.

#### 3.6.3 Case study-3

Tyrosine kinases are enzymes that selectively phosphorylate Tyrosine residues in different protein substrates. In particular, the binding of a signalling molecule with a Receptor Tyrosine Kinase (RTK) activates tyrosine kinase, which is situated in the receptor’s cytoplasmic tail. After this binding, a set of enzymatic reactions happens, which carries the signal to the nucleus, where the kinase regulates protein transcription^50^. The Eph (erythropoietin-producing hepatocyte) receptors are the largest of the RTK families. The cytoplasmic portion of all 14 Eph receptors are highly similar, each containing a membrane-juxtaposing kinase domain, a protein-binding SAM domain followed by a short carboxyl tail PDZ domain-Binding Motif (PBM). The crystal structure of EphA5 in complex with SAMD5 (PDB code: 5ZRZ)^50^, solved at a resolution of 1.89 Å, was analyzed in this study (Figure 9A).

**FIGURE 9.**
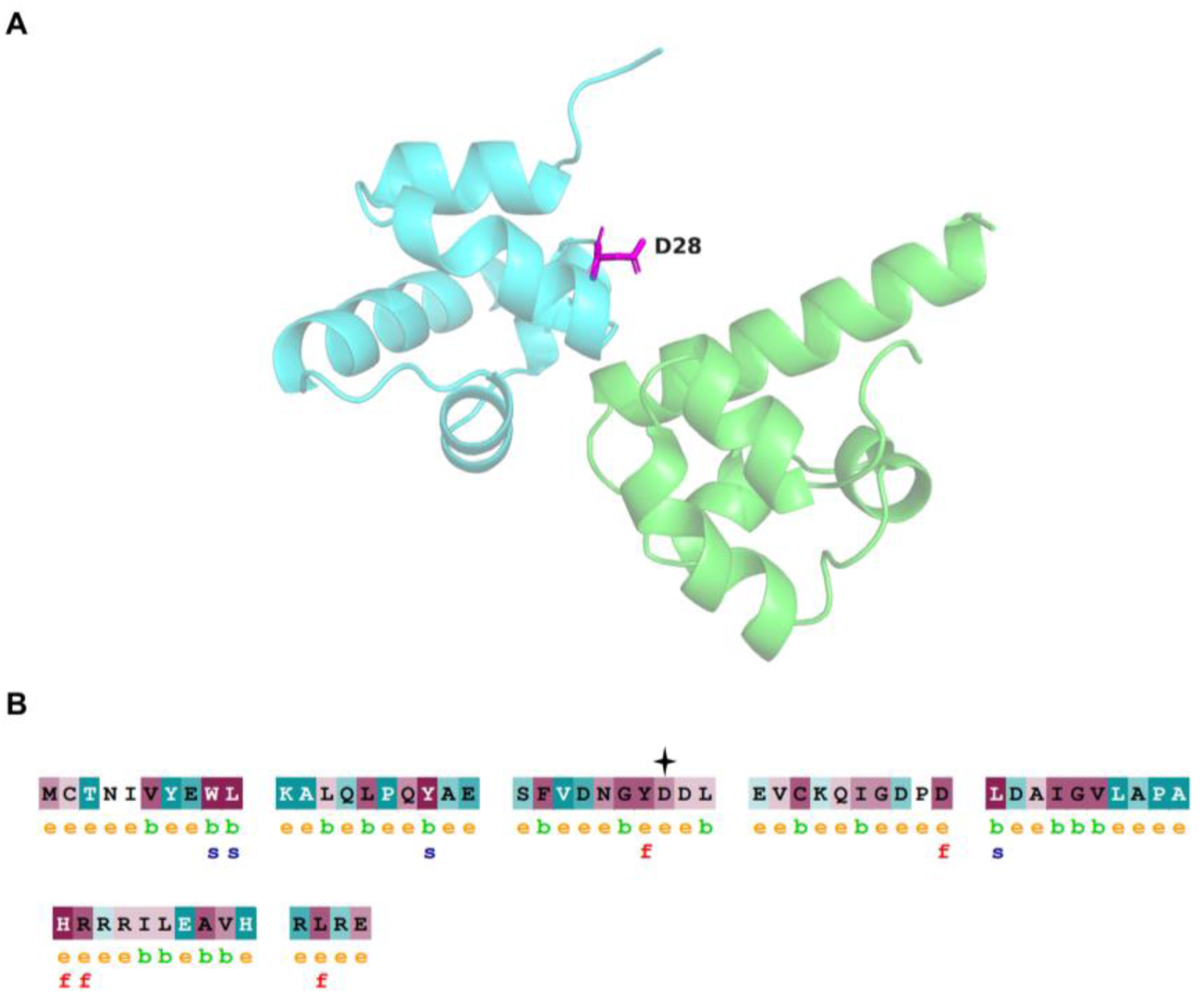
(A) Cartoon representation of EphA5 (green) and SAMD5 (cyan) complex structure (PDB code: 5ZRZ). Asp28 in SAMD5, a residue unique to the rim and experimentally proven to be a hotspot, is shown in magenta sticks. (B) Amino acid sequences colour coded by conservation score for protein SAMD5. The figures were generated using ConSurf^34^. The residues discussed in the case study are marked in black star.

#### 3.6.4 Interface residues unique to distance form crucial inter-protein and intra-protein interactions

#### Met77 of VASH1 and Met66 of VASH2

Met77 of VASH1 in the VASH1-SVBP complex (PDB code: 6J7B) and the equivalent residue Met66 in VASH2 in the VASH2-SVBP complex (PDB code: 6J4P) are both interface residues unique to distance which were computationally predicted to be hotspots. These residues are predicted to be highly conserved (ConSurf grade 9) (Figures 6B and 7B) and contribute favourably to the complex’s overall energy (residue-wise energies -1.66 and -1.39 kJ/mol, respectively). Met66 of VASH2 is a part of the ND crucial for binding with SVBP. Met77 of VASH1, which is part of the predominantly hydrophobic interface patch 1 is experimentally proven to be contributing substantially to the activity of the VASH1-SVBP complex. The experimental mutations M77R+F141R and M77R+V81R+F141R substantially reduced the detyrosination activity^51^. Both Met77 of VASH1 and Met66 of VASH2 interact with Leu 42 of SVBP, which is a residue essential for activity and binding, as proven by experimental mutagenesis studies (Tables 1 and 2). Met77 also forms intra-protein interactions with Trp74, Trp78, Val81, and Phe141, which are residues experimentally proven to be important for binding and activity (Table 1). Interestingly, these intra-protein interactions seem to be conserved in the VASH2-SVBP complex as well (Tables 1 and 2). When an *in silico* mutation of both Met77 of VASH1 and Met66 of VASH2 to all other 19 amino acid types was performed, nearly all the mutations destabilized the respective complexes except for Leu (Figure S9). The ΔΔG value corresponding to the mutation of Met77 of VASH1 to Leu was predicted to be 0.84 kcal/mol, and for other residues, the same lies between 1.99 and 12.45 kcal/mol for other residues. Similarly, mutation of Met66 in VASH2 to Leu predicts a ΔΔG value of 0.39 kcal/mol and ranges between 1.67 and 16.08 kcal/mol for other residues. The primary reason for this destabilization in the mutants is that the hydrophobic interaction network is getting disturbed. This is foreseen, especially since Met77 is part of interface patch1 of the VASH1-SVBP complex, which is predominantly hydrophobic in nature (Figure 6C, left panel). It was also observed that the interaction network of this residue is conserved when Met77 is mutated to Leu, the mutant which did not show much change in the ΔG values (Figure 6C, middle panel). The ΔΔG corresponding to the mutation to Arginine was predicted to be 5.47 kcal/mol. Proximal charged residues could also cause repulsions when mutated to a similarly charged residue causing destabilization in the complex. Figure 6C (right panel) shows that residue 77 mutated to Arg and the nearby positively charged residues, and it is evident that such an Arg residue would not prefer to be placed in a positively charged environment. Moreover, there was a loss of the inter-protein interaction with Leu 42 of SVBP, a residue important for binding and activity, when mutated to any other residue except Val, Ile, Phe, Tyr and Trp. However, in the case of these exceptional residues, there was a loss of intra-protein interactions with some of the residues important for the binding, as described in Tables 1 and 2. Another rationale for the observed effect of *in silico* mutations could be that the amino acid variety seen in the MSA produced by ConSurf showed that apart from Met, Leu is the only residue which is evolutionarily preferred at that position.

**TABLE 1.**
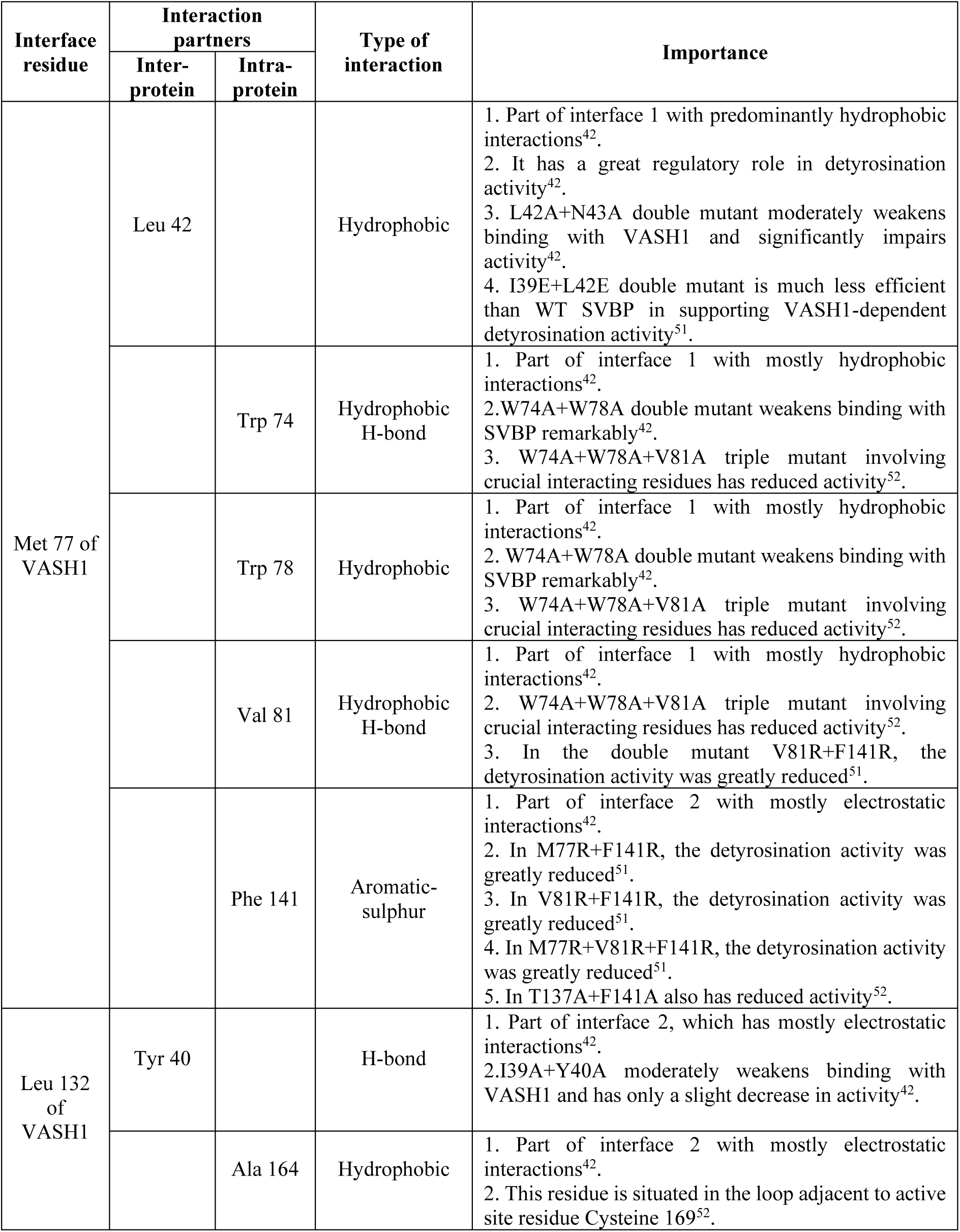
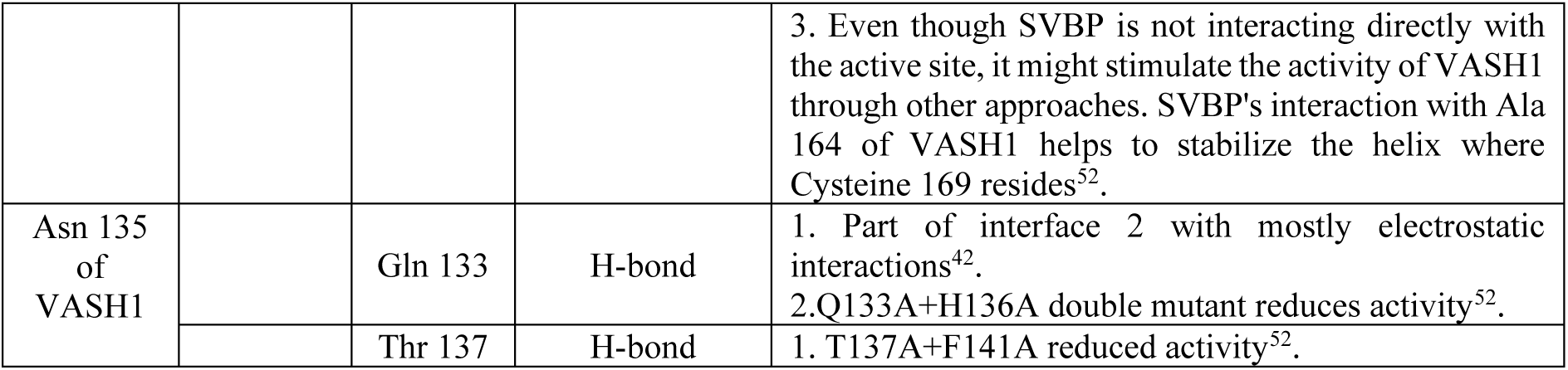
Interface residues unique to distance in VASH1-SVBP complex (PDB code: 6J7B) and some of their important inter-protein and intra-protein partners identified using the all-atom distance cut-off of 4.5Å. The type of interactions were identified using PIC^26^ webserver and the importance of each binding partner are taken from various experimental studies reported in the literature.

**TABLE 2.** Interface residues unique to distance in VASH2-SVBP complex (PDB code: 6J4P) and some of their important inter-protein and intra-protein partners identified using the distance cut-off. The type of interactions and the importance of each binding partner are identified in the same way as in Table 1.

| Interface residue | Interaction partners |  | Type of interaction | Importance |
| --- | --- | --- | --- | --- |
|  | Inter-protein | Intra-protein |  |  |
| Met 66 of VASH2 | Leu 42 |  | Hydrophobic | 1. Interacts with V2c<sup>43</sup>.<br>2. Conserved residue<sup>43</sup>.<br>3.L42A + N43A shows a reduction in complex formation<sup>43</sup>. |
|  |  | Trp 63 | Hydrophobic<br>Aromatic-sulphur<br>H-bond | 1. Interacts with SVBP (Part of ND), conserved<sup>43</sup>.<br>2. W63E + W67E disrupts complex formation, which confirmed that binding with SVBP is essential for activity<sup>43</sup>. |
|  |  | Trp 67 | Hydrophobic | 1. Interacts with SVBP (Part of ND) and conserved<sup>43</sup>.<br>2. W63E + W67E disrupts complex formation, which confirmed that binding with SVBP is essential for activity<sup>43</sup>. |
|  |  | Val 70 | Hydrophobic<br>H-bond | 1. Interacts with SVBP (Part of ND)<sup>43</sup> |
|  |  | Phe 130 | Aromatic-sulphur | 1. Interacts with SVBP (Part of ND)<sup>43</sup> |
| Leu 121<br>of<br>VASH2 | Tyr 40 |  | H-bond | 1. Interacts with V2c and conserved<sup>43</sup>.<br>2. I39A + Y40A showed a reduction in complex formation, which confirmed SVBP binding is essential for activity<sup>43</sup>. |
| Asn 124<br>of<br>VASH2 |  | Gln 122 | H-bond | 1. Interacts with SVBP (Part of ND)<sup>43</sup>. |
|  |  | Thr 126 | H-bond | 1. Interacts with SVBP (Part of ND)<sup>43</sup>. |

It was interesting to observe that while comparing the interaction patterns of the interface residues unique to distance in the two case studies, all the interactions are nearly common across VASH1-SVBP and VASH2-SVBP (Tables 1 and 2). In contrast, interface residues unique to ASA formed interactions that are very specific to each complex, as discussed in the coming sections.

#### 3.6.5 Distance method captures interface residues important for activity which also form a stable interaction network in the unbound and bound forms

#### Thr209 from LmP

Thr209 from LmP of case study-2 is a residue situated in the route to the redox active residue Trp208 of LmP. It is predicted to be an interface residue unique to distance and also a hotspot. This residue has a ConSurf grade of 8 (Figure 8B) and a favourable residue-wise energy of -2.24 kJ/mol. It forms an intra-protein H-bond with Asp211, which is also situated in the route to the radical and forms an ion pair with Arg24 of LmCytC, a crucial ion pair for the electron transfer active complex to form^45^. Earlier experimental studies have reported that the engineered mutants D211N (PDB code: 5AMM)^48^ and D211R (PDB code: 5AL9)^53^ showed reduced activity (Table 3). It also forms an intra-protein interaction with Thr193. It was interesting to observe that both these intra-protein interactions were preserved in both the unbound (PDB code: 3RIV) and bound (PDB code: 4GED) forms of LmP, indicating a stable interaction network formed by Thr209 (Figure 8C). *In silico* mutagenesis to any other residue showed changes in ΔG values in the range of 1.69 and 16.19 kcal/mol, indicating the destabilization of the complex. This was expected as this residue is proximal to the active site and has a regulatory role in the activity. It is of importance that computational approaches are able to identify such important residues in terms of stability and function, which also maintains a stable interaction network from unbound form to bound form.

**TABLE 3.** Interface residues unique to distance and ASA methods in LmP-LmCytC complex (PDB code 4GED) and some of their important inter-protein and intra-protein partners identified using the distance cut-off. The type of interactions and the importance of each binding partner are identified in the same way as in Table 1.

| Interface residue | Intra-protein partner | Type of interaction | Importance |
| --- | --- | --- | --- |
| Thr 209 of LmP | Asp 211 | H-bond | 1. Involved in the most crucial ion pair with Arg 24 of LmCytC, needed for the active electron transfer (ET) complex to form<sup>45</sup>.<br>2. Situated in the route to the redox-active radical Trp 208 of LmP<sup>45</sup>.<br>3. Engineered mutant D211N (PDB code 5AMM) showed only 8% wild-type activity<sup>48</sup>.<br>4. Engineered mutant D211R (PDB code: 5AL9) has reduced activity<sup>53</sup>. |
| Glu 54 of LmP | Asp 50 | H-bond | 1. Part of the secondary binding site<sup>54</sup>.<br>2. D47A+D50A+E54A lowered $K_{cat}$ by 3-fold and the rate of association by 6-fold<sup>48</sup>. |
| Leu 20 of CytC | Arg 16 | H-bond | 1. This residue approaches helix A, which aids in pulling LmCytC away from the ET active site, which is an energetic incentive for LymCytC to move towards helix A<sup>54</sup>. |
|  | Arg 24 | H-bond | 1. Part of the most crucial ion pair with Asp 211 of LmP, which is needed for the proper ET complex to form<sup>45</sup>.<br>2. R24A reduces activity significantly<sup>45</sup>.<br>3. R24A + K98A has same effect as R24A<sup>45</sup>. |

#### 3.6.6 Interface residues unique to ASA form crucial intra-protein interactions Arg44 of SVBP

In the case study of VASH2-SVBP (PDB code: 6J4P), Arg44 of SVBP in the complex is predicted to be an interface residue unique to the rim. This residue is moderately conserved (grade 6) (Figure 7C) and has unfavourable residue-wise energy (2.33 kJ/mol). It forms intra-protein hydrogen bonds with Tyr40 and Leu42, which contribute to the complex formation, as proven by various experimental mutagenesis studies (Table 4). *In silico* mutation to any other residue did not show any change in ΔG except for Pro with a ΔΔG of 3.56 kcal/mol, indicating the destabilization of the complex. While comparing the hydrogen bond network associated with Arg44 in the wild-type and the mutant R44P (Figure 7D left panel and right panel), only three out of six H-bonds are conserved in R44P. Especially, the hydrogen bonding with the conserved residues Tyr 40 and Leu 42 was lost only when mutated to Pro. This interaction was preserved in all other virtual mutation types that resulted in negligible changes in ΔG values. Also, since this residue is situated in a helix, substitution by a Pro residue (“helix breaker”) would not be favourable. Out of all the mutants, the Ala mutant was the complex which showed the least change in ΔG. Interestingly, in this mutant, all the H-bonds in the wild-type were conserved (Figure 7D, middle panel).

**TABLE 4.** Interface residues unique to ASA in VASH2-SVBP complex (PDB code: 6J4P) and some of their important intra-protein partners identified using the distance cut-off. The type of interactions and the importance of each binding partner are identified in the same way as in Table 1.

| Interface residue | Intra-protein partner | Type of interaction | Importance |
| --- | --- | --- | --- |
| Ile 58 of VASH2 | Trp 63 | Hydrophobic | 1. Part of ND and a conserved residue<sup>43</sup>.<br>2. W63E + W67E disrupts complex formation, which confirmed that SVBP binding is essential for the activity<sup>43</sup>. |
|  | Phe 130 | Hydrophobic | 1. Part of ND<sup>43</sup>. |
| Arg 44 of SVBP | Tyr 40 | H-bond | 1. Interacts with V2c and conserved<sup>43</sup>.<br>2. I39A + Y40A reduces complex formation, which confirmed that SVBP binding is essential for activity<sup>43</sup>. |
|  | Leu 42 | H-bond | 1. Interacts with V2c and conserved<sup>43</sup>.<br>2. L42A + N43A showed a reduction in complex formation<sup>43</sup>. |
|  | Met 46 | H-bond | 1. Interacts with V2c<sup>43</sup>. |

#### 3.6.7 Interface residues unique to the rim have a potential role in the recognition, dynamics and specificity of protein-protein complexes

#### Glu54 of LmP

From case study-2, Glu54 of LmP is a less conserved (grade 4) (Figure 8B) interface residue unique to the rim, which has an unfavourable residue-wise energy (4.21 kJ/mol). Even though it does not form any direct interactions with LmCytC, it is reported to be situated in a non-catalytic secondary binding site called helix A, which is vital for complex recognition (Figure 8A). This recognition site is formed by Glu54 along with other negatively charged residues Asp47, Glu49 and Asp50 situated in helix A of LmP. Various molecular dynamics simulation-based studies have reported that LmCytC initially docks to LmP helix A in a non-specific manner and then migrates or crawls towards the ET active site. At the active site, the crucial ion pair between Asp211 of LmP and Arg24 of LmCytC holds the LmCytC in perfect position for the electron transfer^48, 54^. The experimental mutagenesis D47A+D50A+E54A subsided Kcat by 3-fold and the rate of association by 6-fold^48^. Glu54 from LmP is an example of the potential role of interface residues unique to the rim in the recognition and dynamics of the complex. Also, the intra-protein H-bond interactions formed by this residue with Asp50, Met51, and Ile52 are preserved in both unbound (PDB code: 3RIV) and bound forms, indicating the stable interaction network formed by this residue. Except for Pro, which led to a destabilization of the complex (ΔΔG 1.35 kcal/mol), *in silico* mutation to all other residue types had not shown notable effect on the complex stability (ΔΔG ranging between -0.99 and 0.49 kcal/mol). Interestingly the intra-protein hydrogen bond with Asp 50 (Table 3) was conserved in all except Pro. Another intriguing observation was that the *in silico* single mutation to Ala did not destabilize the complex. However, when the same mutation was a part of the triple mutant D47A+D50A+E54A in the experimental studies, it drastically reduced the complex formation. Mainly the interface residues unique to the rim are sensitive as the effect of the mutation in the presence or absence of water is unpredictable. Hence, from this analysis, we suggest that while designing mutagenesis experiments for residues at the periphery of the interface, multiple mutations may be preferred rather than single mutations. In the MSA generated by ConSurf, the amino acid variety percentage of Ala (30.5%) is greater than that of the wild-type Glu (4.9%) at position 54 of LmP. Hence, it is suggested that the classical alanine scanning mutagenesis won’t be ideal for interface residues like this case since the single mutation did not destabilize the complex. Also, a higher occurrence of Ala does not mean that it is always tolerated; hence, evolutionary data is insufficient for designing mutagenesis experiments for residues like Glu54 which are at the periphery of the interface.

Earlier studies have reported that the residues away from the centre of the interface of a protein-protein complex contribute to the specificity of the complex rather than stability^55^. Hence, we probed to identify if residues unique to the rim have a role in the specificity of the complexes. Table 5 shows some of the inter-protein and intra-protein interactions that are specific to either VASH1-SVBP or VASH2-SVBP, which suggests the potential role of interface residues unique to the rim in the specificity of complexes other than recognition and dynamics.

**TABLE 5.** Comparison of interactions in VASH1-SVBP (PDB code: 6J7B) and VASH2-SVBP (PDB code: 6J4P) complexes involving residues that are unique to rim. Inter-protein and intra-protein partners were identified using the distance cut-off. The type of interactions and the importance of each binding partner are identified in the same way as in Table 1.

| Rim Unique | PDB ID | Intra-protein partner | Importance | Inter-protein partner |
| --- | --- | --- | --- | --- |
| Gln 28 of SVBP | 6J4P | Lys 32: H-bond | - | Phe 56: Cation-PI (Lost in 6J7B) |
| Gln 51 of SVBP | 6J4P | Glu 48: H-bond (Lost in 6J7B) | - | Not at interface |
|  |  | Leu 49: H-bond | - | Val 73: Hydrophobic (ND in 6J4P) (Lost in 6J7B) |
|  |  | Phe 54: H-bond (Lost in 6J7B) | - | Tyr 239: Hydrophobic, aro-aro (One of the residues that is involved in shaping the active site of VASH2) (Lost in 6J7B) |
|  |  | Asp 55: H-bond (Lost in 6J7B) | - | Not at interface |
| Gln 29 of SVBP | 6J7B | Lys 25: H-bond (Lost in 6J4P) | - | Not at interface |
|  |  | Ala 27: H-bond (Lost in 6J4P) | - | Not at interface |
| Gln 33 of SVBP | 6J7B | Leu 31: H-bond | - | Pro 68: Hydrophobic (Lost in 6J4P) |
|  |  | Gln 35: H-bond (Lost in 6J4P) | 1.Q35A+R36A impairs binding<sup>42</sup> and disrupts activity<sup>52</sup> and reduces complex formation in 6J4P<sup>43</sup>. | Val 69: H-bond (Lost in 6J4P) |
|  |  | Arg 36: H-bond | 1.Crucial role in maintaining H-bond network in 6J7B<sup>42</sup>.<br>2.Q35A+R36A impairs binding<sup>42</sup> and disrupts activity<sup>52</sup> in 6J7B and reduces complex formation in 6J4P<sup>43</sup>. | Glu 163: H-bond (Int 2 in 6J7B) (Lost in 6J4P) |
| Ile 58 of VASH2 | 6J4P | Phe 56: Hydrophobic | - | Lys 32: Cation-Pi (Lost in 6J7B) |
| Lys 238 of VASH2 | 6J4P | Tyr 239: Cation-Pi (Lost in 6J7B) | 1. Involved in shaping the active site in 6J4P<sup>43</sup>.<br>2. At the interface with SVBP. | Phe 54: Hydrophobic and aromatic-aromatic (Lost in 6J7B) |
|  |  | Leu 240: H-bond (Lost in 6J7B) | - | Met 61: hydrophobic (Lost in 6J7B) |
|  |  |  | - | Phe 57: hydrophobic (lost in 6J7B) |

#### 3.6.8 Interface residues unique to the rim can be hotspots in protein-protein complexes Asp28 from SAMD5

Asp28 of SAMD5 from case study-3 is an interface residue predicted to be unique to the rim according to the chosen criteria. It is also evolutionarily conserved (ConSurf grade 7) (Figure 9B). This residue has an unfavourable residue-wise energy of 4.73 kJ/mol. Yet, it is experimentally validated to be a hotspot residue in the ephA5-SAMD5 complex^50^. In an earlier study, when this residue was experimentally mutated to Ala, it weakened or abolished the binding between EphA5 and SAMD5^50^. Ile58 of VASH2 in the case study of the VASH2-SVBP complex (PDB code: 6J4P) is another residue which is unique to the rim and predicted to be a hotspot at the interface. Hence, this study suggests that while designing mutations for various experiments, one should not only focus on the directly interacting residues. There is a possibility that a residue unique to the rim, which is away from the centre of the interface, could also be a hotspot. Such residues could be crucial for the complex formation as they have a role in the recognition, dynamics and specificity of the complex. Therefore, both distance and ASA methods need to be considered while identifying interface residues; as evident from this study, using only one of them might miss some crucial residues at the interface.

## 4. CONCLUSIONS

The large-scale analyses on the interface residues unique to distance and ASA methods clearly show that an interface residue can form interactions with its partner protein yet not be at the geometrical interface. Conversely, there can be other interface residues located at the geometrical interface yet not forming any interactions with the protein it binds with. The analyses show that the ASA method tends to identify more interface residues as unique to it, and those are predominantly polar in nature. However, the distance method captures more hydrophobic residues, which are highly conserved, contributing favourably to the complex stability and hotspots. From the detailed analysis of specific case studies from the dataset, it was observed that, in general, interface residues unique to the distance method form crucial interactions that hold the proteins together in a complex. Whereas interface residues unique to the ASA method, even though they do not form any direct interactions with the partner protein, they are involved in crucial intra-protein interactions with residues that are critical in complex formation, stability and function. It was also observed that interface residues unique to ASA have an important role in the overall dynamics, recognition, specificity, and function of the complex. Furthermore, the interface residue unique to the rim also could be a hotspot in the complex.

The large-scale interface analysis used to be limited as not very many protein-protein complex structures is known. But, recently developed artificial intelligence-based methods like AlphaFold2 and RoseTTaFold, aids in accurately modelling intricate protein-protein complex structures. Hence, the bottleneck of lack of protein-protein complex structures is being addressed. In these modern approaches, it is also important to understand what exactly plays a pivotal role in forming a protein-protein complex and how to define the interface regions. Overall, observation from this study suggests that distance-based methods are ideal if one is looking for directly interacting residues. However, one should consider a more extensive set of residues identified by both distance and ASA methods as an interface so that the interface residues (other than the directly interacting residues) which have a crucial role in specificity, recognition, dynamics and function are also captured. This comparative analysis would inspire us to provide a common platform in future to use both approaches seamlessly by the user. We hope this study helps better understand the interface of protein-protein complexes and further aid in designing experimental mutagenesis with a wide range of applications in drug design, predicting protein signalling, and 3D modelling of large macromolecular complexes.

## Acknowledgements

The authors thank Dr. Himani Tandon for her input while conceptualizing the project. This research is supported by Parvathy’s PMRF grant and NS’s JC Bose fellowship. Financial support for Parvathy Jayadevan from the Prime Minister’s Research Fellows (PMRF) scheme and the financial support for Yazhini Arangasamy from the Manfred Eigen fellowship from Max Planck Institute of Multidisciplinary Sciences is gratefully acknowledged. Srinivasan Narayanaswamy is a J.C. Bose National Fellow. Sowdhamini Ramanathan is a J.C. Bose National Fellow (JBR/2021/000006) from the Science and Engineering Research Board, India and Bioinformatics Centre Grant funded by the Department of Biotechnology, India (BT/PR40187/BTIS/137/9/2021). RS would also like to thank the Institute of Bioinformatics and Applied Biotechnology for the funding through her Mazumdar-Shaw Chair in Computational Biology (IBAB/MSCB/182/2022).

## Author Contributions

All authors conceptualized the study, and NS and RS coordinated the study. PJ did all the analyses and wrote the first draft of the manuscript. PJ and YA reviewed the results, and YA and RS improved the manuscript.

## Data Availability Statement

PDB codes used in this study can be accessed from RCSB Protein Data Bank.

