## Supplementary Files for "Interfacial residues in protein-protein complexes are in the eyes of the beholder"

### Supplementary text

#### Leu132 of VASH1 and Leu121 of VASH2:

Leu132 from VASH1 in the VASH1-SVBP complex (PDB code: 6J7B) and the equivalent residue Leu121 from VASH2 in the VASH2-SVBP complex (PDB code: 6J4P) of case study-1 are also predicted to be interface residues unique to the distance method according to the chosen criteria. Leu121 of VASH2 is also reported to be a part of the N-terminal domain (ND). Leu132 of VASH1 and Leu121 of VASH2 have ConSurf grades 7 and 8, respectively, indicating high evolutionary conservation (Figures 6B and 7B). Also, their residue-wise energies of -2.264 and -2.71 kJ/mol, respectively, indicate that they contribute favourably to the overall stability of the respective complexes. Both residues form a conserved H-bond with Tyr40 of SVBP, which is important for the complex formation as the experimental double mutant I39A+Y40A moderately weakens binding with VASH1[1] (Table 1), whereas reduced complex formation with VASH2[2] (Table 2). Besides this, Leu132 of VASH1 forms an intra-protein hydrophobic interaction with Ala164, a residue situated in a loop adjacent to the active site Cys169 residue in VASH1. Even though SVBP does not interact directly with the active site, its interaction with Ala164 helps to stabilize the helix where the active site Cys169 resides, aiding SVBP in its allosteric role in regulating enzymatic activity[3]. When both Leu132 of VASH1 and Leu121 of VASH2 were mutated *in silico* to all other 19 residue types, almost all the mutations destabilized the complex except for Met and Phe, which did not show much change. In the case of VASH1-SVBP, the change in $\Delta G$ was observed to be in the range of 1.26 and 4.84 kcal/mol for all mutations except for Met (-0.03 kcal/mol) and Phe (-0.18 kcal/mol). This observation is explained by the disruption of the hydrophobic interactions, specifically the inter-protein H-bond with Tyr40 and the proximal charged residues.

#### Asn135 from VASH1 and Asn124 from VASH2:

Asn135 from VASH1 in the VASH1-SVBP complex (PDB code: 6J7B) is a residue that is a part of the 133QYNH136 motif, which is conserved across VASH1 and VASH2 and is essential in the allosteric role of SVBP in detyrosination activity[1]. It is predicted to be unique to distance according to the criteria that have been chosen, and it is a highly conserved residue with a ConSurf grade 9 (Figure 6B). It also forms interactions that are crucial for the activity of the complex and a favourable residue-wise energy of -1.75 kJ/mol (Table 1). It forms intra-protein interaction with other residues important for the activity of the complex, as described in Table 1. Asn135 is an example of the importance of interface residues unique to distance and their interactions in the functional aspect of the complex. The equivalent residue Asn124 of VASH2 in the VASH2-SVBP complex (PDB code: 6J4P) is also a residue unique to the distance method, with a favourable residue-wise energy of -1.83 kJ/mol. This residue also is evolutionarily conserved, as indicated by its ConSurf grade 9 (Figure 7B). Notably, the intra-protein interactions of Asn135 in VASH1 with some of the residues important for the activity (Table 1) are also conserved in VASH2 (Table 2).

**Leu20 from LmCytC:**

Leu20 of LmCytC of case study-2 is another interface residue unique to the rim that has an indirect role in the dynamics of the LmP-LmCytC complex. As shown in Table 3, this residue is involved in an intra-protein hydrogen bond with Arg 16, a residue that approaches helix A, aiding in pulling LmCytc away from the ET active site, which in turn becomes an energetic incentive for LymCytC moving towards helix A[4] (Table 3). Like most of the residues unique to the rim, this residue is much less conserved (grade 4) and has unfavourable residue-wise energy (0.17 kJ/mol). This residue also has a very stable intra-protein interaction network while comparing the unbound (PDB code: 4DY9) and bound (PDB code: 4GED) forms.

**Ile 58 from VASH2:**

Ile 58, a part of the N-terminal domain of VASH2 in the VASH2-SVBP complex (PDB code: 6J4P), is an interface residue unique to the rim, which is predicted to be a hotspot, less conserved with a ConSurf grade 5 (Figure 7B) and an unfavourable PPCheck energy 1.93 kJ/mol. Even though this residue does not form a direct interaction with any residue in the partner, it forms important intra-protein hydrophobic interaction with residues that are crucial for binding, as validated by experimental mutagenesis (Table 4). Except for Leu, Val, Pro and Met, which do not show much change in the interaction energy, in silico mutation to any other residue results in a positive ΔΔG, indicating a destabilization of the complex. This could be due to the network of hydrophobic interactions getting disrupted or other proximal similar charged residues. The important intra-protein interactions are lost in most of the mutated complexes, and also, according to the residue variety from ConSurf, other hydrophobic residues like Val, Leu and Met are also preferred at this position in the MSA.

**Supplementary figures and tables**

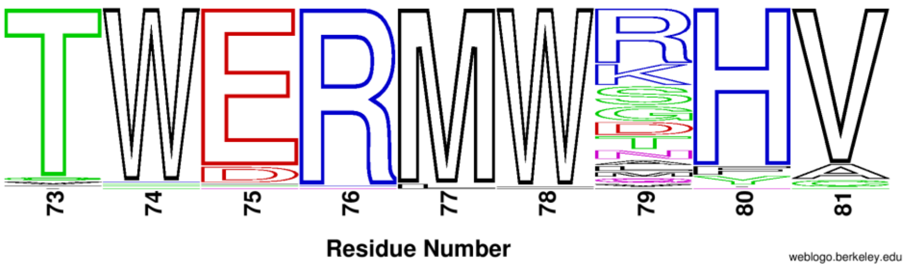


**FIGURE S1** "Amino acid variety" from ConSurf represented in the form of a sequence logo, generated using WebLogo. X-axis represents the residue number, and the size of the logo represents the frequency of occurrence of the specific residue type at that position in the input multiple sequence alignment.

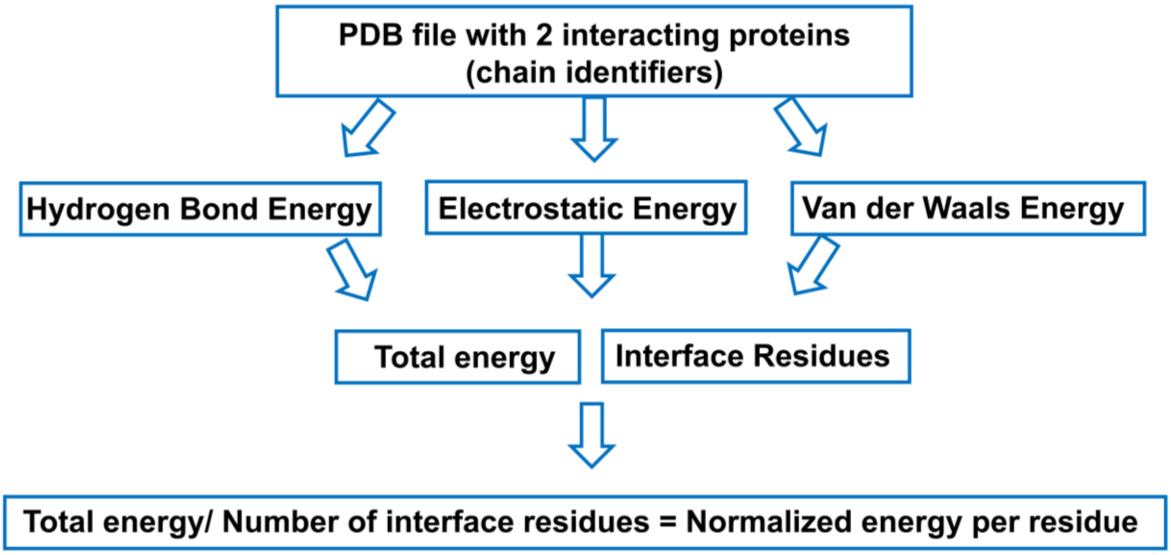


**FIGURE S2** Methodology of PPCheck to calculate inter-residue interaction energies.

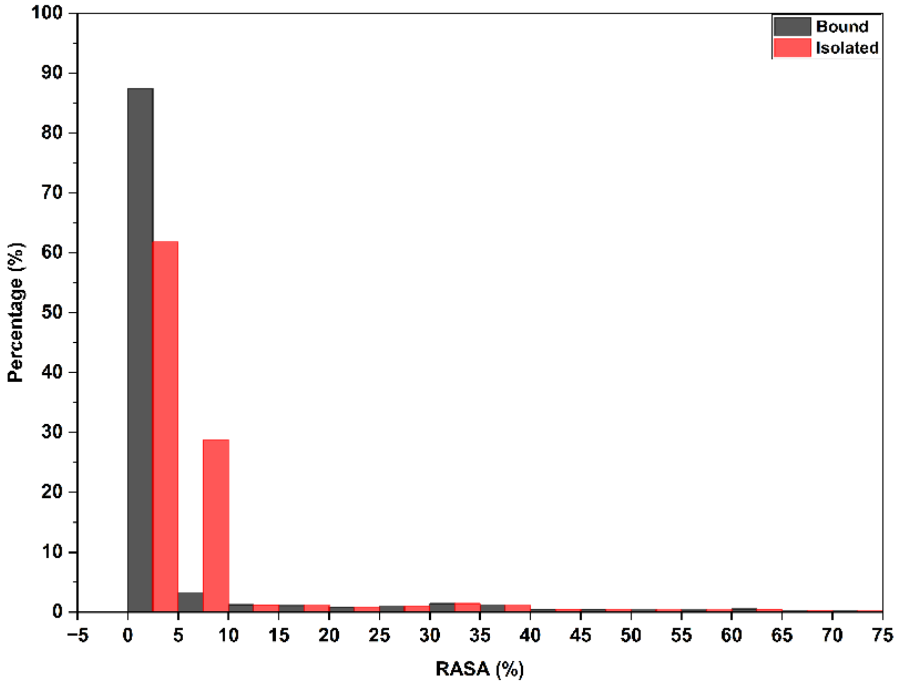


**FIGURE S3.** Histogram showing the percentage of interface residues unique to the distance method with a RASA value belonging to the bins of length 5% ranging from -5 to 75%, both in the bound (grey colour) and the isolated forms (orange colour) of the protein.

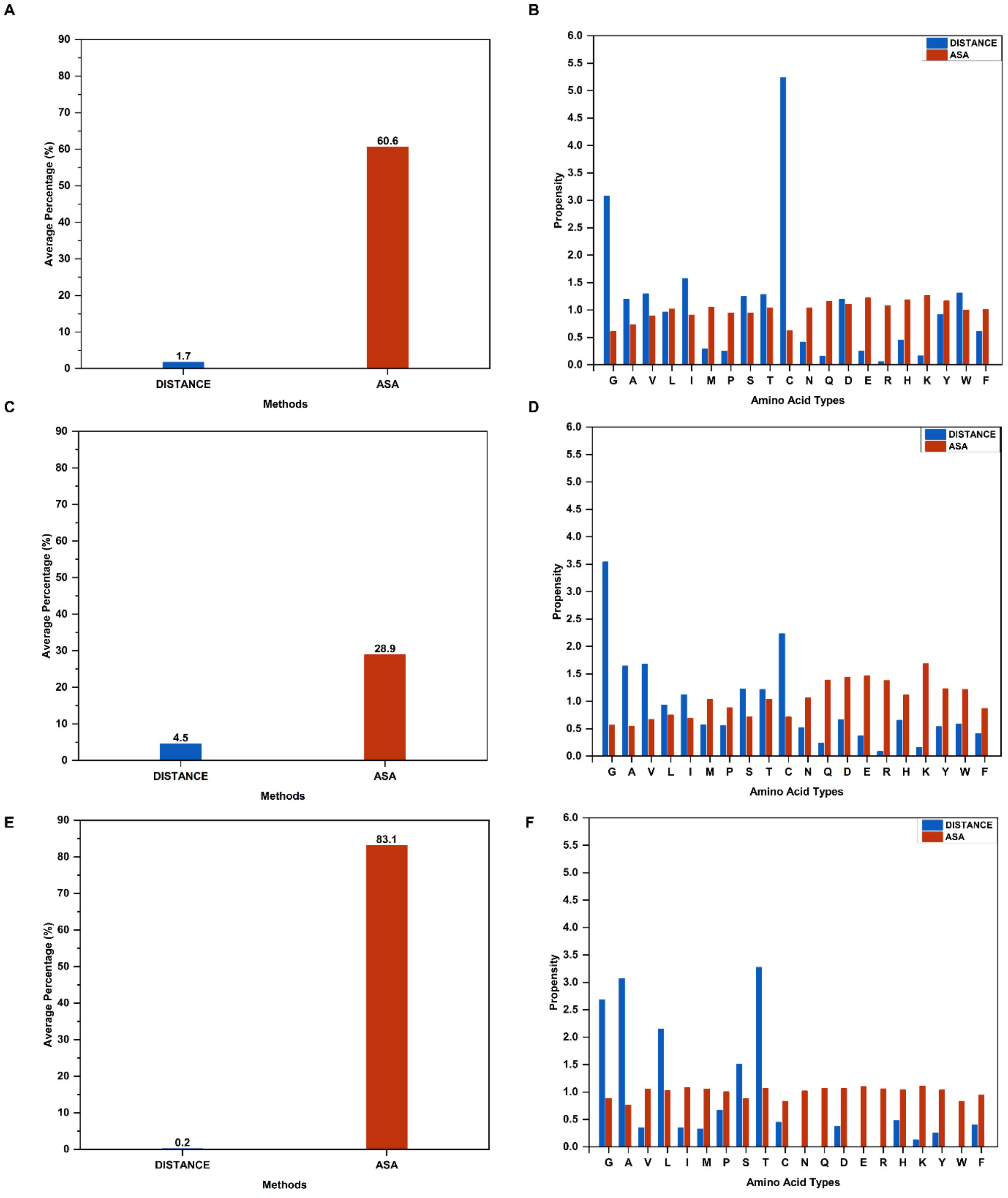


**FIGURE S4** Percentage of interface residues unique to each method for (A) $C^{\alpha}$ – based distance cut-off of 6.5 Å, (C) $C^{\beta}$ – based distance cut-off of 7 Å, and (E) $C^{\beta}$ – based distance cut-off of 4.5 Å. Chau-Fasman propensity calculations for (B) $C^{\alpha}$ – based distance cut-off of 6.5 Å, (D) $C^{\beta}$ – based distance cut-off of 7 Å, and (F) $C^{\beta}$ – based distance cut-off of 4.5 Å.

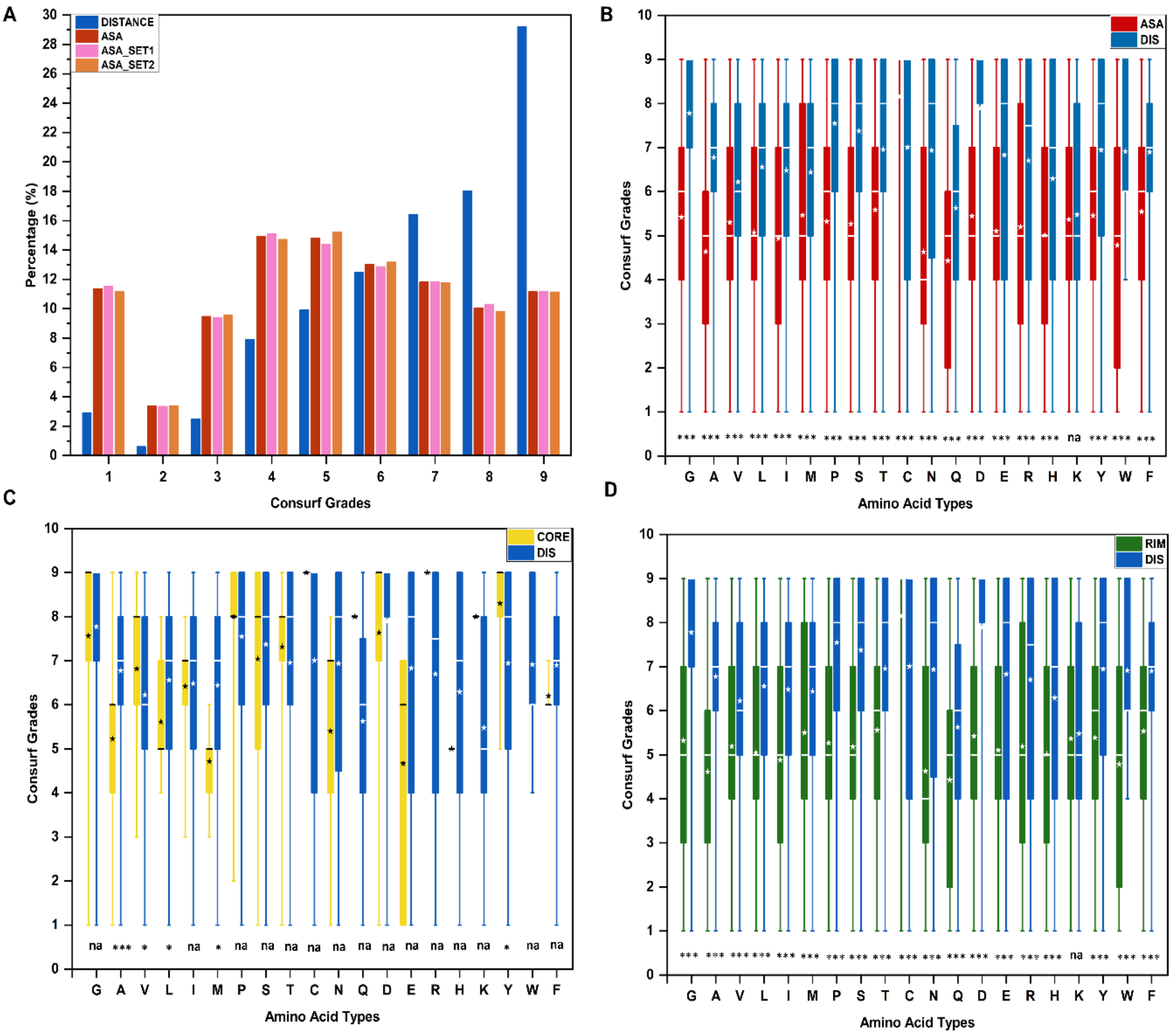


**FIGURE S5** (A) Percentage of interface residues unique to distance method, ASA method and interface residues belonging to ASA_SET1 and ASA_SET2 from random selection for each ConSurf grade from 1-9. Box plot of ConSurf grades for each amino acid residue type for interface residues unique to (B) ASA, (C) core, and (D) rim. "na" indicates either that the distributions are not significantly different or there is not enough data points to do the statistical analysis. A "*" and "***" indicate that the distributions are significantly different, with a p-value in the range of 0.005 - 0.05 and <0.005, respectively. The white star and line on the plot indicate the mean and median of the distribution, respectively.

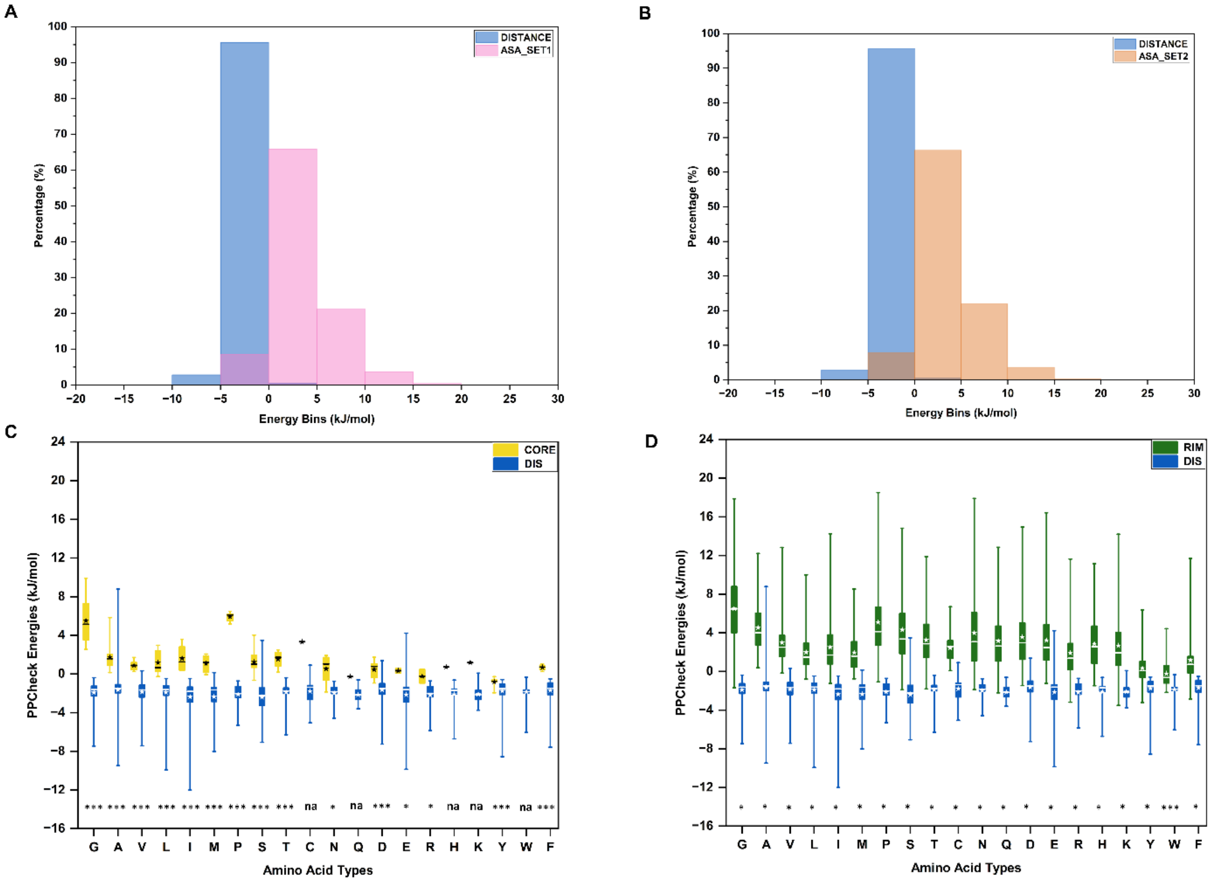


**FIGURE S6** Comparison of the percentage of interface residues unique to the distance method, and the interface residues in (A) ASA_SET1 and (B) ASA_SET2 from random selection, to be in the residue-wise energy bins of length 5 kJ/mol ranging from -20 to 30 kJ/mol. Values beyond 30 kJ/mol are omitted as the values were insignificant. Box plot of residue-wise energies form PPCheck corresponding to each amino acid type for the interface residues unique to distance method, compared to that of (C) core and (D) rim. "na" indicates either that the distributions are not significantly different or there are not enough data points to do the statistical analysis. A "*" and "***" indicate that the distributions are significantly different, with a p-value in the range of 0.005 - 0.05 and <0.005, respectively. The white star and line on the plot indicate the mean and median of the distribution, respectively.

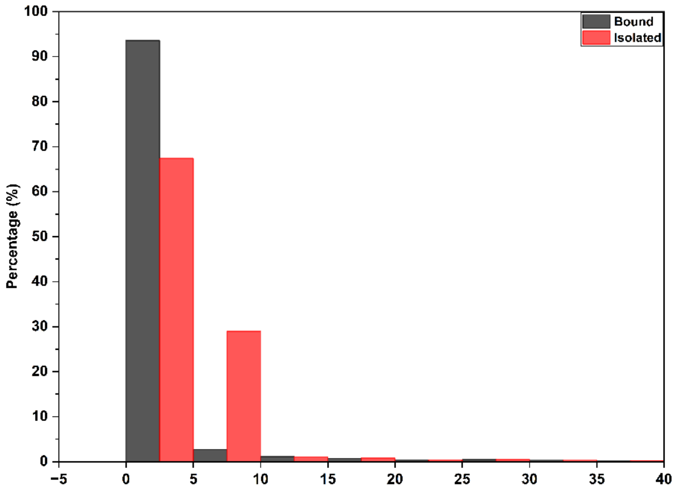


**FIGURE S7.** Histogram showing the percentage of hotspot residues unique to the distance method (secondary shell hotspots) with a RASA value belonging to the bins of length 5% ranging from -5 to 40%, both in the bound (grey colour) and the isolated forms (orange colour) of the protein.

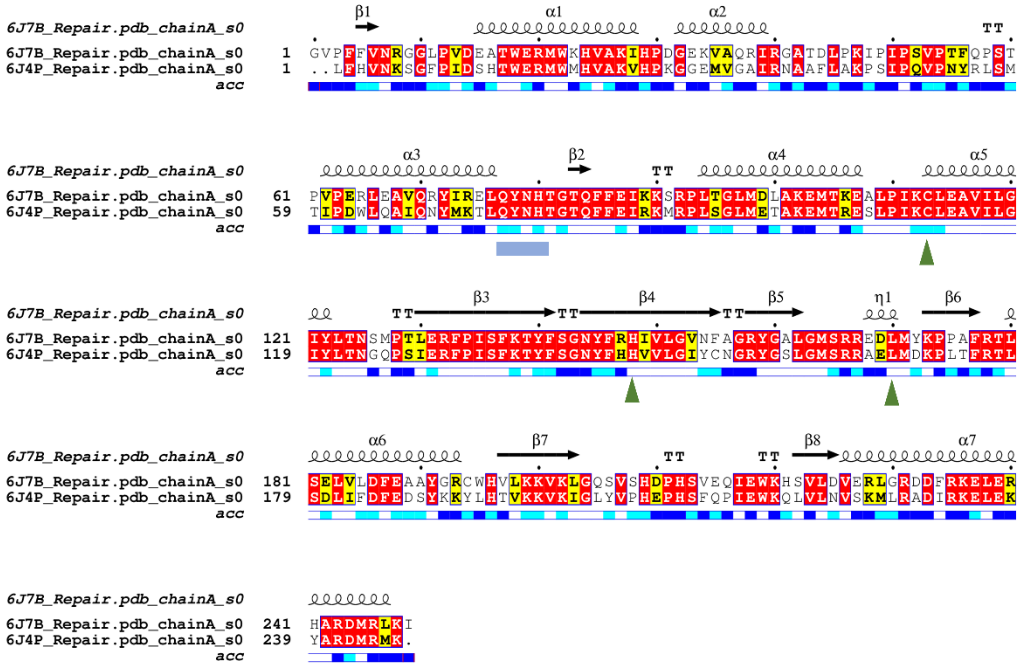


**FIGURE S8** Structure-based sequence alignment of VASH1 and VASH2 using Promals3D. The alignment is rendered using ESPRIPT[54]. The secondary structure elements are annotated at the top of the alignment using the VASH1 structure (PDB code: 6J7B). Corresponding accessibility values are shown at the bottom. Blue, cyan and white indicate solvent accessible, partially accessible and inaccessible regions, respectively. Red and yellow coloums in the alignment show fully conserved and conservatively substituted residues, respectively. The catalytic triad and 133QYNH136 motif conserved between VASH1 and VASH2 are shown in green triangles and blue rectangles, respectively.

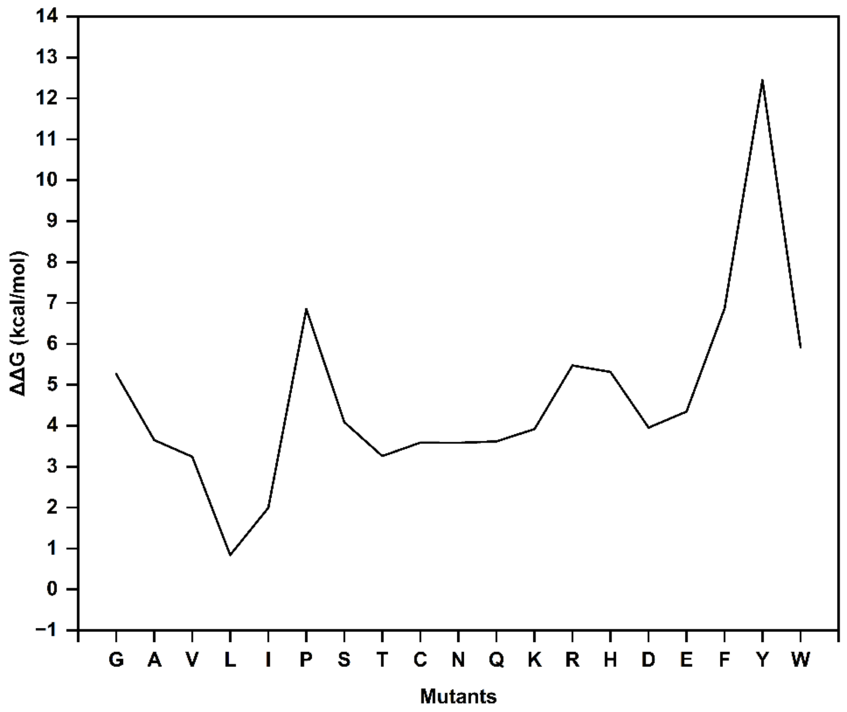


**FIGURE S9** Line plot showing ΔΔG values upon in-silico mutagenesis of Methionine 77 of VASH1 from VASH1-SVBP complex (PDB code 6J7B) to all other 19 residue types. The *in silico* mutagenesis was performed using the PositionScan module of FoldX. A positive ΔΔG value indicates that the mutation destabilized the complex, and a negative value indicates the stabilization of the complex.

**TABLE S1** Relative accessible surface area (RASA) values of the interface residues unique to distance in each case study analysed. RASA values are from naccess and are given for both the isolated and complexed forms of the protein. All the residues unique to distance have an RASA value $< 7\%$ in both forms indicating that these residues are buried in both forms.

| UNIQUE TO DIS | PDB ID | RASA VALUE IN THE UNBOUND FORM (%) | RASA VALUE IN THE BOUND FORM (%) |
|---|---|---|---|
| Methionine 77 of VASH1 | 6J7B | 6.3 | 0.5 |
| Leucine 132 of VASH1 | 6J7B | 5.4 | 0.2 |
| Asparagine 135 of VASH1 | 6J7B | 3 | 2.8 |
| Threonine 209 of LmP | 4GED | 3.1 | 2.1 |
| Met 66 of VASH2 | 6J4P | 6.3 | 0.5 |
| Leucine 121 of VASH2 | 6J4P | 4.8 | 0.5 |
| Asparagine 124 of VASH2 | 6J4P | 2.1 | 1.9 |
| Phenylalanine 130 of VASH2 | 6J4P | 6 | 0.8 |
